# Survey of nucleotide-specific Rab GTPase interactions reveals multiple Rab effectors

**DOI:** 10.64898/2026.09.15.751902

**Authors:** Tabitha A. Peterson, Christopher P. Ptak, Mark A Stamnes, Robert C. Piper

## Abstract

Rab GTPases control myriad molecular functions by acting as a nucleotide-dependent switch for protein-protein interactions localized in discrete subcellular regions (Zerial and McBride, 2001). Finding additional molecular roles for Rab GTPases depends on expanding the identification of their nucleotide-dependent interactions. Here we compare the interactome of Rab GTPases using a comprehensive large-scale yeast 2-hybrid approach powered by quantitative high-throughput sequencing, which was used to find interactions of the major mammalian Rab isoforms locked in a GDP or GTP conformation. These data revealed an expanded set of interactions specific for GTP-bound Rab proteins, with many known Rab effectors capable of binding a wider repertoire of Rab partners than previously appreciated. We also identified the RGS domain of SNX13 and SNX14 as a new Rab GTPase-binding domain that may bridge these ER-localized proteins with Rab5 and Rab11 endosomal compartments, respectively.

## INTRODUCTION

Heterotrimeric and small GTPases share a common core architecture and GTP-binding domain. Both these enzymes alter their conformation when bound to GTP to promote binding to downstream effectors, some of which accelerate hydrolysis of GTP via a GAP (GTPase-accelerating protein) activity (Siderovski and Willard, 2005; Bos et al., 2007; Cherfils and Zeghouf, 2013). Rab GTPases comprise a large subfamily within the collection of small molecular weight GTPases (∼25 kDa) consisting of 6 central β-sheets coordinated by 5 α-helices that provide the well conserved guanine-nucleotide binding fold (Vetter and Wittinghofer, 2001) also present in heterotrimeric G proteins. Binding of GDP or GTP induces conformational changes in the ‘switch’ regions to allow different repertoires of protein interaction while at the same time regulating their association with GDI and the exposure of covalently attached geranylgeranyl lipid tails to cellular membranes into which they are inserted (Rak et al., 2003; Ullrich et al., 1993). Over 60 Rab proteins in humans serve multiple functions throughout the cell by engaging effectors on specific membrane compartments (Hutagalung and Novick, 2011). A key feature of Rab GTPases is their differential localization across cellular compartments, which localizes their activity and the protein interactions they regulate to specific cellular regions (Seabra and Wasmeier, 2004; Pfeffer, 2013).

A major requirement for understanding the full function of Rab proteins is knowing their various nucleotide-specific protein:protein interactions. Some Rab proteins have been found to have multiple partners that likely engage under different cellular states and locations, emphasizing the need to identify specific Rab-interacting proteins and where they may occur. Still other Rab GTPases remain with a paucity of identified interactors leaving their function and how they perform that function unclear. The complexity of Rab interactions and activities is increased by the recognition that each Rab protein may have multiple effectors and that each effector may have capacity to work with multiple Rab proteins. Much work on Rab proteins encompasses both broad and focused screens for Rab interacting proteins using a variety of approaches. This has led to identifying structurally defined domains that mediate Rab interaction for nucleotide exchange activity or effector/GAP activity including DENN, bMERB, RILPHD, TBC/RABGAP, RUN, PH, and GRIP domains (Mott and Owen, 2015; Rai et al., 2016; Marat et al., 2011; Callebaut et al., 2001; Barr, 1999). The defining feature of Rab GTPases is that nucleotide-dependent conformational changes enable them to engage distinct sets of binding partners. A subset of these partners, which control GTP exchange and hydrolysis, regulate the Rab cycle itself. More broadly, the interactions most relevant to Rab function are those that depend on nucleotide state, as these proteins fall under Rab regulation. Identifying such nucleotide-specific partners therefore provides a direct route to defining the cellular roles of individual Rab GTPases.

Several large-scale interaction screens have been used to define protein interaction networks across the proteome, including those involving Rab GTPases. Yet high-throughput generic approaches do not resolve the nucleotide-dependent interactions that underlie Rab function because they do not distinguish between GDP- and GTP-bound states. Grafting mutations originally identified in the Ras GTPase allows Rab proteins to be constrained in their GTP-bound (Q61L in Ras) or GDP-bound (S34N in Ras) conformations. These mutations have long provided tools to probe the function and interactions of Rab proteins within distinct GTP (“active”) or GDP (“inactive”) states (Der et al., 1986; Feig and Cooper, 1988; Li and Stahl, 1993; Stenmark et al., 1994). Similarly, these conformationally locked mutants have been used extensively in biochemical and genetic approaches, including yeast 2-hybrid (Y2H) assays, to identify interacting proteins (Fukuda, 2010; Kail et al., 2008; Langemeyer et al., 2012).

We previously developed DEEPN (Dynamic Enrichment for the Evaluation of Protein Networks), which performs Y2H selections in batch liquid culture under controlled stringency (Pashkova et al., 2016). Deep sequencing of prey-encoding plasmids enables comprehensive and quantitative comparison of interaction profiles across different baits, ensuring equivalent library representation and allowing direct assessment of specificity (Peterson et al., 2018; Krishnamani et al., 2018).

Mapping of prey–vector junctions permits determination of reading frame and interacting fragments, facilitating rapid identification of interaction domains within complex libraries (Krishnamani et al., 2018). One of the missing ingredients in these analyses has been the availability of high-quality Y2H libraries that provide improved representation across the proteome, are not biased by differential mRNA abundance in the cells or tissues from which libraries are derived, and are enriched in ORF-encoding DNA fragments rather than the 3′ untranslated regions that plague older Y2H libraries generated from cDNA synthesis.

Here, we generated a high ORF-density library from the human ORFeome and used it to systematically screen mouse Rab GTPases, producing a comparative dataset that identifies shared and differential nucleotide-conformation-sensitive interactions across the mouse Rab GTPase family.

## METHODS

### Materials: Antibodies, Plasmids, and Strains

Polyclonal anti-myc antibodies were purchased from QED Biosciences (cat#18826). Goat anti-rabbit 680LT antibodies were from Licor (cat#925-68021). Monoclonal anti-HA antibody (Biolegend cat#901514), monoclonal anti-GFP (Santa Cruz cat#sc-9996), and polyclonal anti-mCherry (EMD Millipore cat#AB356482) were used as indicated. Plasmids and where they were used in this study are listed in **Supplementary Table 1**. Yeast strains were PJ69-4A MATa (James et al., 1996) which housed the bait plasmids and PLY5725 MATα (*gal4*Δ::KanR *trp1*Δ::LoxP *gal80*::LoxP, *his3*Δ1 *leu2*Δ0 *lys2*Δ0 *ura3*Δ0), which housed the prey plasmids such as the human ORFeome library.

### Plasmid Construction

The bait plasmids, p*TEF1*-GBD and p*Tef1*\*-GBD, contained the Gal4 DNA binding domain, *TRP1*, and KanR. Rab ORFs were designed using yeast codon bias, and the C-terminal CAAX box was replaced with AAAA to prevent prenylation-dependent membrane association. Prey plasmids were made using the *LEU2*- and AmpR-containing plasmid pGal4-AD by insertion between SfiI sites downstream of the Gal4 activation domain.

### Human ORFeome Library Construction

Human ORFeome 8.1 plasmid DNA pools (Transomic Technologies, Huntsville, AL) were amplified by PCR using NEBNext polymerase (cat#: M0541, New England Biolabs, Ipswich, MA) and a biotinylated primer recognizing the Gateway vector backbone (/5Biosg/TTYTTATAATGCCAACTTTGTACAARAAAG). Purified products were pooled according to the initial complexity of the ORFeome pool from which they were amplified. Amplicons were fragmented using dsDNA Fragmentase (NEB cat#M0348S) according to the manufacturer’s guidelines for 30 s, 1 min, and 1 min 30 s, with reactions halted by addition of EDTA to 10 mM, to generate a range of fragment sizes. Biotinylated fragments corresponding to flanking regions of the amplicons were removed by subtraction over Avidin-Agarose beads (Sigma-Aldrich cat#A9207).

The remaining fragments were ligated to Y-oligo adaptors, allowing ligation into SfiI-cut pGal4-AD after amplification with oligos 5′-TAATCGATAGGCCTCCCG and 5′-AGGCTAGCCGGCCCAGCC and digestion with SfiI. A separate set of fragments was amplified with oligos 5′-GAGTGGTGGCAACTCTGT-GGCCGGCCCAGCCGGCCATGT and 5′-TCCCGTCTTCTATTATTCATCTA-GGCCTCCCGGGCCATGT to generate ends compatible with Gibson assembly. This parallel Gibson strategy was used to preserve rare inserts containing internal SfiI sites that would otherwise be destroyed during restriction cloning. Ligation and assembly products were transformed, grown, and maxi-prepped in batch (Invitrogen cat#K210017), with 1% of each pool plated to estimate the number of independent bacterial plasmid clones generated.

To generate the Human ORFeome Y2H_TAB yeast library, the yeast strain PLY5725 was transformed with the maxi-prepped Human ORFeome fragment DNA library using the LiAc method and plated onto CSM-Leu-Met to select for transformants. After 2 days of growth at 30 °C, colonies were harvested, pooled, resuspended in YPD containing 30% glycerol, frozen, and stored in aliquots at −80 °C. This procedure generated a dense ORF fragment domain-mapping library from amplified and fragmented human ORFs. As a comparison Y2H library, the normalized universal mouse cDNA Mate & Plate library purchased from Clontech Laboratories (cat#630483) was moved into PLY5725 yeast using the same high-efficiency transformation procedure.

### DEEPN (Dynamic Enrichment for Evaluation of Protein Networks)

DEEPN screens were conducted across a panel of Mouse Rab GTPases in their GTP and GDP bound conformations as originally described, with modifications reported more recently (Pashkova et al., 2016; Peterson et al., 2018; Krishnamani et al., 2018). Briefly, Rab GTPase bait plasmids in p*TEF1*-GBD or p*Tef1*\* were transformed into PJ69-4A (*MATa trp1-901 leu2-3,112 ura3-52 his3-200 gal4Δ gal80Δ LYS2::GAL1-HIS3 GAL2-ADE2 met2::GAL7-lacZ*) and mated with PLY5725 (*MATα his3Δ trp1Δ leu2Δ ura3Δ lys2Δ gal80Δ gal4Δ*) cells harboring the Human ORFeome Y2H_TAB prey library in pGal4-AD. GDP and GTP-locked Rab bait pairs were screened in parallel alongside p*TEF1*-GBD vector-alone controls in each run. Each bait-containing population was grown as a matched population and then split into nonselective (+His) and selective (-His) conditions, allowing direct comparison of prey representation before and after Y2H selection. Selective growth was performed without 3-aminotriazole after verifying that bait constructs did not self-activate when co-expressed with the pGal4-AD vector (**Supplementary Figure 1A**).

After growth, prey library fragments were recovered by PCR and subjected to 2 × 150 paired end sequencing on an Illumina HiSeq4000. Sequencing yielded an average of approximately 3.5 million reads per sample, of which approximately 95% mapped to hg38. Reads were mapped to hg38 using MAPster and processed through the DEEPN analysis workflow. Mapped reads were converted to gene level counts through exon matching, while junction spanning reads were analyzed by BLAST to identify prey gene fragments, determine whether fragments were in the correct translational reading frame, and define fragment boundaries. These data were integrated with read depth profiles across annotated transcripts to map interacting regions within candidate prey proteins and were collated for each interaction.

Candidate interactions were identified using the StatMaker module, requiring agreement between two independent statistical estimates: a previously used Bayesian posterior probability of enrichment (P) > 0.8 and a DESeq2 log2 fold-change over vector corresponding to > 3-fold enrichment (p ≤ 0.01), together with a positive Bayesian-adjusted enrichment estimate (AdjEnr) over vector and a percent threshold where junctions bridging the coding region of interest were ≥ 80% in the proper reading frame (e.g., in-frame, forward). All DESeq2 analyses used poscounts size-factor normalization, which substitutes 0.5 for zero counts before computing geometric means and retains genes absent from some samples. Although the DEEPN framework can support three-way comparisons among vector, GDP-locked, and GTP-locked baits, interactions were operationally called using pairwise comparisons, generating interactor sets independently for GDP and GTP state baits. This analysis was designed to identify interactions above significance thresholds rather than infer relative interaction strength between nucleotide states, as empirical prey rankings were less informative than subsequent binary Y2H matrix validation assays used to assess interaction specificity and strength.

To identify Human ORFeome library prey plasmids that enrich nonspecifically on the p*TEF1*-GBD vector independent of any bait fusion (e.g., ‘garbage’ interactors), we performed two DESeq2-based analyses on the existing Rab GTPase Y2H dataset. First, we tested each of 17 independent Vector-Selected/Vector-Non-Select replicate pairs (shared as negative controls across the 36 Rab bait experiments) for genes significantly enriched in Vector-Selected vs Vector-Non-Select conditions with no bait present (DESeq2, p ≤ 0.01 along with positive fold change). Prey that met these enrichment criteria in 7 of 17 independent vector samples were retained as vector background interactors. Second, to capture interactors obscured by competition in large cultures, 2mL aliquots of the vector-alone transformant population were inoculated into 75 mL selective medium and grown to generate 20 ‘bottleneck’ cultures. These were pooled, sequenced, and compared to four different pairs of non-select independent vector replicates as the reference condition. Vector-only interactors were identified by applying an in-frame:forward junction-quality floor of ≥85%, an enrichment requirement of ≥2.5-fold on either the raw within-culture selected/non-select ppm ratio or the Bayesian model-free Bait-vs-Vector enrichment score (Enr1), and a DESeq2 p-value < 0.1 for the Bait-vs-Vector comparison. Prey plasmids meeting these criteria in at least 2 of the 4 independent reference-vector comparisons were retained as additional vector-alone interactors, together producing an initial nonspecific interactor set of 70 genes. Prey with a UniProt-annotated subcellular localization of predominantly secreted (n = 6), or whose interacting fragment mapped to the extracellular or lumenal face of an integral membrane protein by its annotated transmembrane topology (n = 13), were also excluded as inaccessible to a cytosolically-expressed bait. Nonspecific interactors were defined not just by gene identity but at the level of the specific interacting fragment, and this fragment-level resolution allowed us to retain genuine, bait-specific interactions even for genes whose other fragments were flagged as nonspecific. As a final step of curation, the interacting fragment recovered with each Rab bait was compared directly against each of the 17 vector-alone datasets. A fragment enriched in any single vector-alone set that overlapped the fragment recovered with the bait was flagged as nonspecific (**Supplementary Table 2)**. The full set of Rab interacting fragments (**Supplementary Table 3)** and nonspecific Vector interacting fragments with descriptions of vector junctions, statistics, and read-depth profiles are available at: https://doi.org/10.6084/m9.figshare.28678313

#### DEEPN_26

DEEPN, MAPster, and StatMaker were updated to run natively on Apple Silicon (ARM64) Macintosh computers using the DEEPN software previously reported (Krishnamani et al., 2018). All three were ported from Python 2/PyQt4 to Python 3/PyQt5 and distributed as macOS application bundles requiring no separate runtime installation. Porting was carried out with the assistance of Claude Code (Anthropic), and a detailed description of the software is available as a supplement: DEEPN_Software_Description at: https://doi.org/10.6084/m9.figshare.28678313

The underlying analytical methods were unchanged from Krishnamani et al. (2018). Raw reads are aligned to hg38 using HISAT2, counts are generated per gene by matching reads to annotated exon coordinates, and BLASTn identifies candidate prey junction fragments and characterizes them by reading frame and position. StatMaker’s statistical model was updated to a DESeq2 approach adapted from Velasquez-Zapata et al. (2021), added alongside the prior JAGS framework. The original JAGS-based Bayesian model, which the analyses reported here rely on, was preserved unchanged for continuity with the previously validated analysis; DESeq2 is offered as a complementary approach. In both, variance is estimated from duplicate vector-alone control conditions run in parallel under nonselective and selective growth conditions, as before. Hit-calling was further refined to require agreement between the DESeq2 fold-change and the Bayesian-adjusted enrichment estimate before a gene is called, and to report two complementary p-values — computed directly (p-value_raw) or normalized for baits that strongly deplete the non-select population (p-value_norm) — so users can choose the more appropriate one for a given screen.

FragFinder is a new module for reconstructing prey plasmid inserts from screen data. Junction positions mark the nucleotide coordinates where the Gal4 activation domain sequence meets an inserted library fragment, and these positions are displayed along the reference sequence of the gene of interest. For a selected junction, FragFinder rescans the dataset’s raw BLAST and junction files for every read supporting that exact junction, reapplying the same hit-quality filter used during initial processing, and returns the longest such read to define the insert boundary. Read-depth coverage across the gene is plotted separately from the mapped .sam files.

MAPster is available as a standalone application and is also launchable from within DEEPN, so alignment, count generation, and junction extraction can run as a single workflow. MAPster also supports exporting alignments as .bam files via a bundled copy of samtools, for tools that require BAM input. DEEPN’s own pipeline uses .sam files directly.

Software releases:

DEEPN: https://github.com/RobertPiper/DEEPN/releases/tag/v7
MAPster: https://github.com/RobertPiper/MAPster/releases/tag/v4
StatMaker (standalone): https://github.com/RobertPiper/DEEPN/releases/tag/statmaker-v7

### Pull-down binding experiments

GST fusion proteins were produced in BL21(DE3) bacteria, purified over Glutathione Sepharose 4B, eluted with 25 mM reduced glutathione in PBS, dialyzed three times against PBS, and stored at 4 °C. For each binding reaction, 200 μg of each GST protein was immobilized onto 50 μL total Glutathione Sepharose 4B packed bead volume. GST-protein-bound beads and bacterial cell lysates were incubated by gentle rotation in a total volume of 500 μL PBS containing 0.1 mg/mL BSA, 0.1 mg/mL casein, and 0.02% Triton X-100 for 1 h at 22 °C. Beads were washed three times with 4 °C PBS containing 0.02% Triton X-100, eluted with 100 μL 4× Laemmli sample buffer containing 2-mercaptoethanol, and heated for 5 min at 70 °C. Samples were subjected to SDS-PAGE followed by immunoblot analysis.

### NMR spectroscopy

For NMR spectroscopy, bacteria expressing ^15^N GST-TEV-SNX14 RGS were grown to an OD600 of 0.7. Cells were resuspended in minimal medium containing 1× M9 medium, 1 g ^15^N ammonium chloride, 0.1 mM calcium chloride, 1 mM magnesium sulfate, 3× vitamins (MEM vitamin solution), 1 μg/mL thiamine HCl, 4 g/L glucose, and 10% ^15^N Celtone (Cambridge Isotope Laboratories, Tewksbury, MA), and grown for an additional hour at 37 °C prior to induction with 0.5 mM IPTG. Samples underwent initial purification as described above. After cleavage of the GST tag with TEV protease, ^15^N SNX14 RGS and Rab11 Q70L were purified by gel filtration over Superdex 75.

Samples were dialyzed into 50 mM Na₂PO₄, 20 mM NaCl, 2 mM MgCl₂, 1 mM DTT, 1 mM GTP, pH 6.95. Two-dimensional ^1^H-^15^N HSQC spectra were acquired using 50 μM ^15^N SNX14 RGS with or without 50 μM Rab11 Q70L in the same buffer containing 10% D₂O. Spectra were acquired on a 800 MHz Bruker Avance II NMR spectrometer at 25 °C. Data were processed using NMRPipe (Delaglio et al., 1995), and spectral contour plots were generated using POKY (Lee et al., 2021).

### Cell culture and transfection

COS-7 cells were maintained at 37°C with 5% CO_2_ in Dulbecco’s modified Eagle’s medium (Gibco) supplemented with 10% fetal bovine serum (Gibco) and 50 U/mL penicillin, 50 μg/mL streptomycin. Plasmid transfections were carried out using Lipofectamine LTX Reagent with Plus Reagent (Invitrogen, Cat# A12621).

### Model building

The MICAL-L1 bMERB model was made by superposing the AlphaFold model of human MICAL-L1 (UniProt Q8N3F8; Jumper et al., 2021; Varadi et al., 2024) onto the MICAL1 bMERB domain in a complex with Rab10 at Site 1 and Site 2 (PDB 5LPN) (Rai et al., 2016). The SNX14 and SNX13 RGS domain models (AlphaFold; UniProt Q9Y5W7 and Q9Y5W8; Jumper et al., 2021; Varadi et al., 2024) were superposed onto RGS8 in the RGS8–Gαi3 crystal structure (PDB 2ODE; Soundararajan et al., 2008), and GTP-bound Rab11 (PDB 2HV8; Eathiraj et al., 2006) and Rab5 (PDB 1R2Q; Terzyan et al., 2004) were superposed onto the Gαi3 G domain.

### Microscopy

COS-7 cells were seeded into glass-bottom dishes (MatTek, P35G-1.5-20-C). At 18 h post transfection, images were captured using a Leica SP8 confocal microscope. Image processing included acquisition of Z-stacks and conversion of the four center stacks into maximum intensity projections using LAS-X software.

### Co-immunoprecipitation

At 18 h post transfection, cells were washed once with DPBS and treated with 1 mL Versene at 37 °C and 5% CO₂ for 5 min. Cells were transferred to a 1.5 mL Eppendorf tube and pelleted at 500 × g for 5 min at room temperature. Supernatant was gently removed, and pellets were resuspended in 500 μL ice-cold lysis buffer containing 1× PBS, 1× EDTA-free protease inhibitors (Roche), 2 mM Pefabloc, 1% DDM, and 0.2% CHS. Lysates were incubated on ice for 30 min and clarified by centrifugation at 12,000 rpm for 20 min at 4 °C. Supernatants were incubated with 17.5 μL GFP nanobead volume per reaction at 4 °C with rotation for 1 h. Beads were washed three times with PBS containing 0.011% DDM and 0.002% CHS, and proteins were eluted with 2× Laemmli sample buffer for analysis by western blot. Equivalent starting material was analyzed in parallel to assess pulldown efficiency.

### Western Blot Yeast Protein Expression

Yeast cells were resuspended in 450 μL of 0.2 N NaOH for 5 min at 25 °C, repelleted, and solubilized in 75 μL of 8 M urea, 5% SDS, and 10 mM Tris, pH 6.8. Samples were analyzed by SDS-PAGE, immunoblotting, and imaging using either a Licor imager or FluoroChem 8800 (Alpha Innotech).

### Binary 2-hybrid Assays

Cells expressing bait and prey fusion proteins were mated on YPD plates overnight, streaked onto CSM-Leu-Trp to select for diploids, and spotted onto CSM-Leu-Trp and CSM-Leu-Trp-His plates. Bait and prey fusion protein expression was verified before mating. Baseline production of His3 from bait plasmids in PJ69-4A/PLY5725 diploids was assessed by monitoring growth on CSM+His and CSM-His plates incubated at 30 °C for 3 days. Interaction strength was assessed semiquantitatively by serial dilution endpoint growth, based on the number of dilutions that produced opaque growth on selective medium. Prey fragments were tested in matrices across multiple Rab bait proteins to assess Rab specificity and nucleotide-state specificity.

### DEEPN screen quality control

Variability across non-select Y2H populations was assessed by comparing each Rab bait population to an empirical null model derived from p*TEF1*-GBD vector-only bait populations. Gene abundances (ppm) were converted to counts, normalized for sequencing depth using median-of-ratios size factors, and gene-wise dispersion parameters were estimated using a negative binomial model implemented in DESeq2. For each gene, a Pearson-type residual statistic was calculated to measure deviation of observed prey abundance from vector-derived expectations while accounting for read depth, gene-specific overdispersion, and minimum sampling noise for low-count observations. Sample deviation was summarized as the median absolute residual across genes, providing a measure of the typical magnitude of departure from the vector-defined null.

Expected baseline variability was established by leave-one-out (LOO) cross-validation of vector populations, in which each vector replicate was excluded in turn and scored against a model rebuilt from the remaining vectors. The resulting LOO score distribution defined the empirical background, including the maximum observed deviation. Rab bait populations were then scored against the full vector model to quantify bait-dependent distortion of prey representation under nonselective growth conditions. For bait constructs that impaired yeast growth, expression was reduced using a weaker *TEF1* promoter (p*Tef1*\*-GBD), and these samples were additionally evaluated using a within-cohort LOO model constructed from the reduced-expression populations themselves, providing a scale-matched null for downstream analyses. For populations under selection for positive Y2H interaction (e.g., –His), where enrichment concentrates reads into a relatively small number of prey genes, median residual statistics were no longer informative. Instead, these samples were assessed using a Pearson correlation coefficient (PCC) between normalized prey abundances and a reference expectation derived from vector-only populations grown under the corresponding selective conditions. This metric quantifies the extent of preservation versus distortion of the expected prey distribution under selective pressure. Correlations were computed on the raw normalized abundance scale, which is particularly sensitive to bait-driven redistribution among dominant prey species that is otherwise lost under rank-based measures (e.g., Spearman correlation). Full methods and R-scripts used for quality control calculations are available as a DEEPN_QC supplement at: https://doi.org/10.6084/m9.figshare.28678313

## RESULTS

### Construction of a High-Density Yeast Two-Hybrid Library for DEEPN Analysis

The general principle of the yeast 2-hybrid (Y2H) system is to identify interacting proteins through the reconstitution of a transcriptional activator when two fusion proteins associate. A key advantage of this approach is that it employs defined proteins to detect binary interactions. While intermediary proteins may occasionally contribute, a substantial proportion of detected interactions are direct. The use of protein fragments, rather than exclusively full-length proteins, represents an additional strength of Y2H. Fragments can delineate regions responsible for binding and may facilitate interaction by liberating domains that are otherwise occluded within complex tertiary structures or regulated by intramolecular interactions.

Applying Y2H to comprehensively compare interactomes is more challenging. Traditional formats rely on defined matrices of bait and prey plasmids, limiting scalability. Although large-scale Y2H studies have generated extensive datasets, they typically assess binary interactions among full-length proteins and do not capture interactions mediated by protein fragments or conformational variants. Consequently, interactions involving proteins such as Rab GTPases, whose binding partners depend on nucleotide-specific conformations, can go undetected.

Our previous studies demonstrated the utility of performing Y2H screens in batch mode combined with high-throughput DNA sequencing to monitor enrichment of library plasmids under conditions requiring a positive Y2H interaction. This approach, termed Dynamic Enrichment for the Evaluation of Protein Networks (DEEPN), enables quantitative comparison of interactomes across multiple conditions. By employing relatively permissive selection, DEEPN captures transient interactions that would otherwise be obscured under highly stringent yeast 2-hybrid selection, where competitive exclusion leads to clonal dominance by a limited number of high-affinity interactions. Finally, enriched prey plasmids can be computationally analyzed to identify specific coding sequence subregions responsible for binding. When deployed with a library containing a high density of ORF fragments, DEEPN provides detailed information on the domains mediating interaction. Moreover, tracking fragments that fail to enrich can indicate regions that are insufficient for binding. By quantifying each human ORF library element before and after selection, interactomes can be compared across multiple baits, provided that ample and consistent sampling is maintained.

To enhance the DEEPN approach, we engineered a new Y2H library containing fragments derived from the Human ORFeome v8.1 (**Figure 1A**). This collection comprises approximately 17,000 sequence-verified open reading frames (ORFs) and represents a broad swath of proteins expressed in human cells. While not fully comprehensive, it provides extensive coverage of the human proteome, and future iterations may incorporate alternative ORF collections to expand this approach. ORFs were amplified from ORFeome subpools and ligated into a simplified *LEU2*-containing prey plasmid downstream of the Gal4 activation domain (Gal4-AD). Before ligation, amplicons were enzymatically fragmented to avoid sequence bias associated with mechanical shearing. Unfragmented amplicons and terminal fragments containing vector-derived ends were removed using avidin-coated beads. The remaining fragments were then ligated into the prey vector to generate a high-complexity library enriched in coding sequences. A similar strategy was previously used to construct a yeast genomic fragment library that achieved high fragment density per ORF, facilitated by the fact that approximately 70% of the yeast genome is protein-coding (Peterson, et al., 2018). Likewise, amplification from the human ORFeome enriched the resulting ORFeome library for coding sequences. The plasmid prey library was transformed into PLY5725 yeast (*his3Δ trp1Δ leu2Δ ura3Δ gal80Δ gal4Δ*), expanded, and aliquoted, allowing the same prey population to be used across multiple Rab GTPase baits.

**Figure 1.**
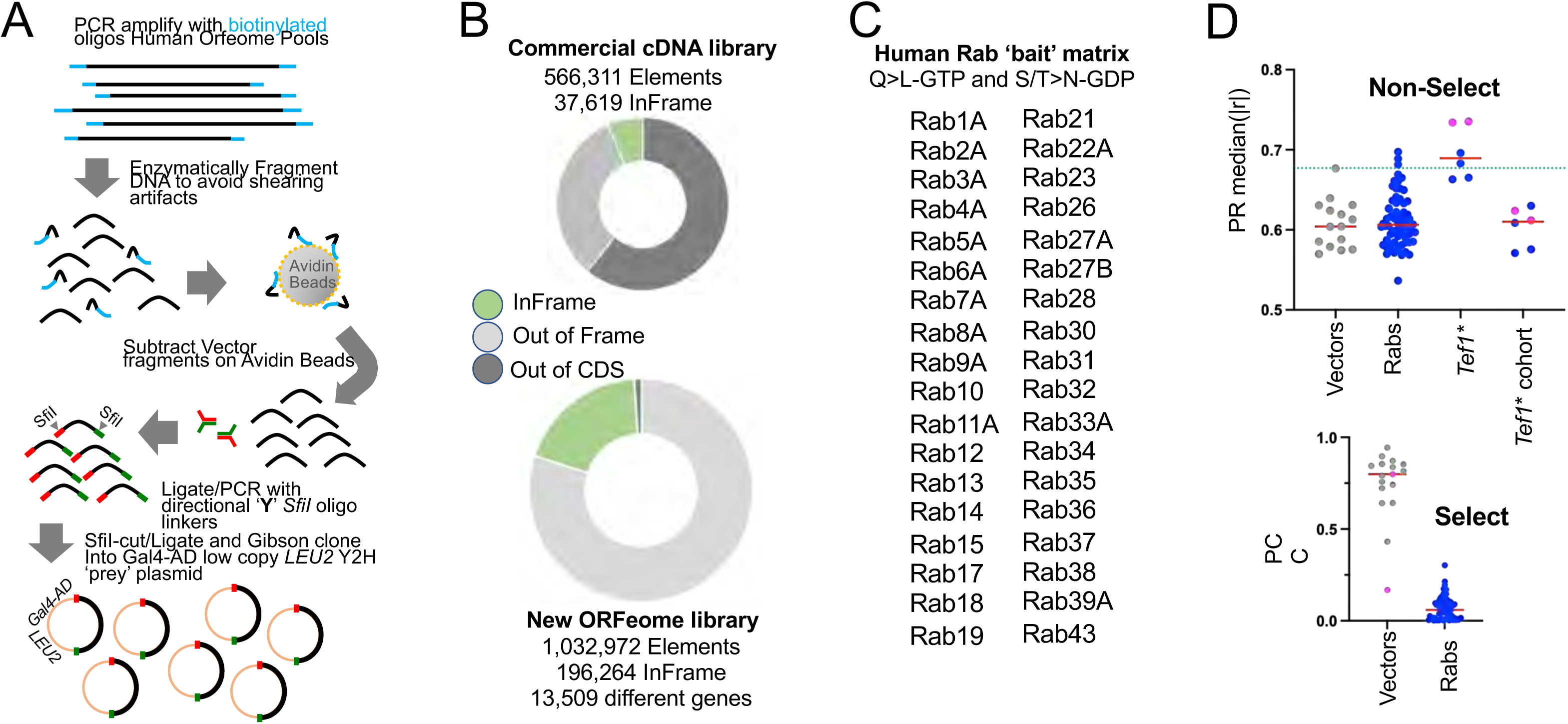
Generation of human ORFeome fragment library. **A.** Inserts from the human ORFeome v8.1 plasmid pool were amplified with biotinylated oligos and enzymatically fragmented to avoid shearing artifacts. After depleting vector-containing ends over avidin beads, fragments were conjugated and amplified with directional ‘Y’ linkers containing SfiI sites and ligated into a low-copy *LEU2* plasmid downstream of the Gal4 Activation Domain. **B.** Comparison of a commercial Y2H cDNA library from Clontech and the human ORFeome fragment library. ‘Prey’ plasmids from libraries housed in transformed yeast populations were amplified and sequenced using an Illumina paired-end workflow. Individual fragments/elements were identified by their unique junctions (aka, 5′ junctions) that bridge the connection of the Gal4 transcriptional activation domain and ORF fragments. Analysis of these junctions determined the translational frame of each insert. The commercial Y2H library contained only 37,619 in-frame elements. Most of the elements contained 3′ untranslated fragments outside of the coding sequence. The new ORFeome fragment library had 196,264 different fragments that were in frame with the Gal4 Activation Domain. **C.** List of Rab proteins used in comparative DEEPN screens with the human ORFeome library. Each Rab GTPase was engineered as a GTP-locked conformation (Q>L) or GDP-locked conformation (S/T>N), comprising 72 total Rab GTPase ‘bait’ plasmids. **D.** Quality analysis of Y2H populations under conditions that do (Select) or do not (Non-Select) select for positive interactions supporting growth in media lacking histidine. (Top) Per-sample median(|r|), where r is the depth-aware Pearson residual from a DESeq2 intercept-only negative-binomial model. Vector controls (p*TEF1*-GBD empty-vector populations, n=15) scored under leave-one-out (LOO) cross-validation define baseline compositional variability (Vectors); the dashed green line marks the LOO maximum at 0.677. The Rab bait populations (Rabs) were scored against the full p*TEF1*-GBD vector reference and fell within this envelope (mean = 0.61, SD = 0.03; 93% within vector mean ± 2 SD). Populations driven by the weak *Tef1*\* promoter are shown two ways: (1) scored against the p*TEF1*-GBD vector reference (p*Tef1*\*-GBD vectors in pink, p*Tef1*\*-Rabs in blue), with all 6 breaching the vector envelope (0.66–0.74); and (2) grouped into their own LOO cohort (p*Tef1*\* cohort) and clustering with a variability in line with the other samples. (Bottom) Pearson correlation coefficient (PCC) of each sample’s count profile against the vector reference. Vector controls (*pTEF1*-GBD empty-vector populations, n=15) were scored under leave-one-out (LOO) cross-validation and maintain high correlation under selection (PCC = 0.43–0.94). The two p*Tef1*\*-GBD vector populations (pink) are shown for comparison. Rab bait populations (Rabs) were scored against the full p*TEF1*-GBD vector reference and collapse to PCC ≤ 0.30 (median = 0.054), indicating that bait-specific prey enrichment during selection drives each population away from a vector-alone profile.

High-throughput DNA sequencing of plasmid junctions spanning the fusion between the Gal4-AD and the inserted library fragment revealed that the yeast-housed library contained approximately 1.03 million unique elements, of which ∼19% encoded in-frame coding sequence (CDS) fragments (**Figure 1B**). Plasmid-carrying yeast populations were propagated in the absence of Y2H selection (+His) and analyzed by high-throughput sequencing of prey inserts isolated by PCR. Unique plasmid elements were identified by sequencing reads spanning the junction at the 3′ end of the Gal4-AD. Because fragments can insert in either orientation and across three reading frames, approximately one-sixth of inserts are expected to be in-frame. For comparison, a commercially available Y2H library generated from normalized cDNAs derived from multiple tissues exhibited a total complexity of approximately 0.57 million elements, with only 6.7% encoding in-frame CDS fragments. Although those inserts were directionally cloned, the library was heavily dominated by sequences corresponding only to 3′ untranslated regions due to oligo-dT priming, rendering a substantial portion of the library unsuitable for interaction screening. The DEEPN software workflow maps reads against 19,117 annotated genes. Sequencing of 15 independent p*TEF1*-GBD vector-alone control populations (53.1M reads total) revealed that 13,509 of these genes were represented in the Y2H human ORFeome library.

### Data Processing

Our original JAGS Bayesian model (Pashkova, et al., 2016) estimates the probability of authentic enrichment on one bait over two others, enabling 3-way comparisons. One application for Rab GTPases is finding prey enriched on the GTP-locked form over both the GDP-locked form and vector alone. However, pairwise comparisons proved sufficient to identify interactions specific to each nucleotide state, without requiring direct comparison of enrichment levels between baits. Enrichment was treated as binary such that prey either met the significance threshold or not, and the resulting interactor lists were compared. This avoided any need to interpret that relative enrichment strength was meaningful across conditions, which would only require the same competing interactor pools were present under each bait.

Each bait’s enrichment was assessed against vector independently, but both baits were included jointly in a single 3 way model for each run. The model estimates a baseline abundance and a general selection response for each gene, shared across every bait and vector in a given run rather than specific to any one of them. Including a second bait’s data gives the model an additional independent read on that shared baseline for each gene, reducing noise that would otherwise be attributed to any single bait’s apparent enrichment. The 3 way comparison was therefore run even when only one bait’s enrichment over vector was of interest. The DESeq2 model was chosen for the improved DEEPN software because of its widespread use, open source implementation, and prior validation on DEEPN data, with variance estimated from in-run vector conditions as described in Methods (Velásquez-Zapata et al., 2021).

Prey plasmids enriched under selective conditions on vector alone defined the nonspecific interactor set. Enrichment of this set was reproducible across multiple runs, with good agreement in both nonselective and selective conditions (**Figure 1D**). But strong vector-only interactors could potentially dominate the enrichment space available to all library plasmids and obscure weaker vector-only enrichments in a large population. To broaden the detection of vector-only interactors, we analyzed 20 smaller vector-only cultures, creating a bottleneck that allowed otherwise diluted library elements to better compete for enrichment. The final nonspecific interactor list is the union of all vector-alone selective populations, large and small, accumulated across multiple runs (**Supplementary Table 2**). This list defines these interactors at the fragment level, not just the gene level. This allowed enrichment of the same gene on a bait of interest to still be detected and favorably scored as a candidate provided that enrichment mapped to a different fragment. Overall specificity was further supported by the structure of the dataset. Each Rab GTPase was screened in both GDP- and GTP-locked states, and most interactions were observed in only one nucleotide state. This means the same Rab in the opposite nucleotide state serves as a built-in negative control, independent of any within-run statistical test. Moreover, an interaction that is consistently enriched across multiple replicates of one bait but absent across dozens of other Rab datasets provides a cross-experiment specificity argument that no single-run statistical test can match.

### Rab-GTPase screening

**Figure 1C** shows the 36 different human GTPases used as ‘bait’ plasmids for this study with the results of their overall screen shown in **Tables 1, 2 and Supplementary Table 3.** Each GTPase was expressed as either the GTP-bound conformation with a Leu substitution at a conserved glutamine corresponding to Ras Q61L (Der et al., 1986) or Asp substitution at a conserved Ser/Thr residue (S31 in HRas; Feig and Cooper, 1988). In addition, the CAAX prenylation motif was mutated to AAAA. Most were expressed from the high-level constitutive *TEF1* promoter, and all were verified as expressing full-length protein at comparable levels *via* immunoblotting for in-frame epitope and did not produce a positive Y2H interaction when combined with an empty prey plasmid (**Supplementary Figure 1A,B**). Expression of any of the ‘bait’ Rab-GTPases had no effect on growth except for Rab1 and Rab18.

**Table 1.** Rab- and GTP-specific interactors. Prey recovered as interactors of a single Rab GTPase exclusively in its GTP-locked conformation. Each column lists prey genes for the indicated Rab bait. 141 interactions across 32 Rab GTPases, drawn from the filtered dataset of 522 Rab-prey interactions (**Supplementary Table 3**) after removal of nonspecific and topologically inaccessible prey (**Supplementary Table 2**). Previous interactions described in BioGRID are in blue.

| Rab19 (1/6) | Rab19 (2/6) | Rab19 (3/6) | Rab19 (4/6) | Rab19 (5/6) | Rab19 (6/6) |
| --- | --- | --- | --- | --- | --- |
| ALDH1A3 | CKMT1B | IQSEC2 | NTNG1 | RAB36 | TOX |
| APBB1 | DCDC5 | KBTBD3 | OSBPL2 | RAB3A | TPMT |
| ARID5A | DDX1 | L2HGDH | OSGIN2 | RGS7 | TTL |
| C5orf34 | DVL3 | LHX4 | OXA1L | RNF157 | TTLL4 |
| C7orf31 | EML1 | MSTO1 | PAF1 | SH3BP4 | ZCCHC7 |
| C9orf152 | GRHL1 | MYCBPAP | PIAS1 | SYT11 | ZGRF1 |
| CKMT1A | HSPBAP1 | NEFL | POLR2H | THUMPD2 | ZNF670 |

| Rab38 | Rab22 | Rab9 | Rab36 | Rab2 | Rab8 | Rab27B | Rab31 |
| --- | --- | --- | --- | --- | --- | --- | --- |
| ALS2CR12 | ABHD3 | FANCC | ADAD1 | CCDC186 | EPB41 | NUP54 | HMGXB4 |
| BCAS2 | BEST3 | GIMAP4 | BAG6 | CES1 | HEXDC | TRAF4 | MTRF1L |
| DNAJB13 | GABRB1 | HPS4 | DAB2IP | FAM217A | KRT3 | TRIP6 | PBLD |
| INTU | HECTD2 | PSMD12 | ERICH1 | FUS | ZBTB48 | UFL1 | PLD2 |
| KRT14 | NFE2L2 | PTPRA | NOS1AP | TTK | ZNF713 |  |  |
| KRT23 | PRKRA | RMND5B | SMYD1 |  |  |  |  |
| KRT33B | RBSN | ZC3H8 |  |  |  |  |  |
| KRT34 | SMAP1 |  |  |  |  |  |  |
| KRT9 | TLE1 |  |  |  |  |  |  |
| LMNB2 |  |  |  |  |  |  |  |
| NDUFV2 |  |  |  |  |  |  |  |
| TRIM17 |  |  |  |  |  |  |  |

| Rab37 | Rab43 | Rab14 | Rab26 | Rab28 | Rab30 | Rab5A | Rab11 |
| --- | --- | --- | --- | --- | --- | --- | --- |
| CTDSPL2 | CBWD5 | DDX20 | KRT27 | CAMK4 | GDPD5 | ASMT | LARS2 |
| IFIH1 | DNAJB2 | POC1B | SIL1 | EFCAB13 | TSHZ1 | ZNF567 | OR8D2 |
| KIAA0226L | EEF2K | SNX29 | TMSB15B | IRF4 | ZNF582 |  |  |
| ZNF680 | USP33 |  |  |  |  |  |  |

| Rab12 | Rab15 | Rab23 | Rab27A | Rab32 | Rab33A | Rab34 | Rab35 |
| --- | --- | --- | --- | --- | --- | --- | --- |
| ATXN10 | CREB3L4 | DNAJC7 | CAPN6 | DARS | NF2 | MSH2 | FBXO7 |
| SLC12A4 | GK5 | PPP1R15A | STRIP2 | KRT31 | SYBU | SLC37A3 | LRRC3B |

| Rab3A | Rab4A | Rab6A | Rab7A | Rab13 | Rab18 | Rab21 |
| --- | --- | --- | --- | --- | --- | --- |
| PRICKLE3 | FAM184A | GTPBP4 | UTP3 | ZFAND3 | TBC1D9B | MFSD9 |

**Table 2.** Interactors recovered with two or more Rab GTPases. Prey genes (rows) enriched on two or more of the 36 Rab baits (columns), each divided into GTP-locked and GDP-locked sub-columns. Filled cells indicate a recovered interaction; black fill denotes interactions previously documented in BioGRID; red fill denotes novel interactions identified in this study.

To ensure that downstream Y2H interaction profiles can be compared directly across baits we confirmed that all nonselective libraries contain equivalent prey plasmid composition prior to selection (**Figure 1D**). This analysis tests whether each bait is exposed to the same underlying library, such that differences observed after selection reflect bait-specific interactions rather than differences in prey availability.

Datasets were filtered to the 13,509 genes represented in the library using the union of all non-select populations. To assess whether individual bait populations distorted prey abundance beyond baseline library variation, each non-select library was scored against a vector-derived null model built from p*TEF1*-GBD control populations (n = 15) using a DESeq2-based negative binomial framework, with gene-specific dispersion used to calculate Pearson-type residuals, and sample deviation summarized as the median absolute residual across genes (median_abs_r). Vector control populations exhibited low variability by LOO analysis (**Figure 1D**; median_abs_r: median = 0.6042, 95th percentile = 0.6506, maximum = 0.6766). Unselected Rab-bait yeast populations fell within or close to this empirical background range when scored against the vector-derived reference constructed from the full set of p*TEF1*-GBD vector-alone samples, indicating that prey library composition and relative abundance were largely preserved across populations. These results show that prey-library yeast populations before selection remain highly comparable across different baits and support valid quantitative comparison of interaction profiles following selection.

Bait constructs with Rab1 and Rab18 produced a pronounced growth defect in yeast under non Y2H-selective conditions (+His). Rab1 and Rab18 share high levels of identity with Ypt1 (71% and 51%, respectively) and Sec4 (55% and 48%, respectively), both essential yeast Rab GTPases and high expression as ‘activated’ Rab-GBD fusion proteins likely perturbs these essential functions. Impaired growth can distort the starting prey library composition by selecting for plasmids that confer a growth advantage independent of a bait–prey Y2H interaction, for example through multi-copy suppression, which in turn would invalidate cross comparison with other Rab Y2H populations. To mitigate this effect, these baits were expressed from a reduced-strength promoter vector (p*Tef1*\*-GBD), which alleviated the growth defect. Pearson residual analysis of this cohort (that included both vector and Rab baits under the *Tef1*\* promoter) showed that prey library composition and median_abs_r values were slightly elevated out of the range of the standard (p*TEF1*-GBD) vector distribution. When evaluated as a separate cohort using the LOO approach (n = 6, two p*Tef1*\*-GBD vector controls plus the 4 reduced-expression Rab bait samples), scores fell within a narrow band (median_abs_r: median = 0.6103, 95th percentile = 0.6285, maximum = 0.6301), consistent with strong internal consistency within this subset (**Figure 1D**). Downstream enrichment analyses for these baits were therefore performed against matched reduced-expression p*Tef1*\*-GBD vector controls so that interaction signals are interpreted relative to the appropriate baseline.

Following Y2H selection, 12,388 genes were retained after union filtering across the full dataset. Because selection concentrates signal into a relatively small number of strongly enriched prey, median absolute residual statistics lose power to resolve differences among post-selection populations. We therefore summarized selected samples using the Pearson correlation coefficient (PCC) between normalized counts and the vector-derived reference expectation (**Figure 1D**, bottom). Bait-specific prey enrichment was evident as a marked reduction in correlation to the aggregate vector reference. Replicates of p*TEF1*-GBD vector-alone samples showed Pearson coefficients spanning 0.43–0.95 by LOO, whereas Rab-bait populations fell to 0.00–0.20, indicating substantial bait-driven redistribution of prey composition under selective conditions. Together, these results show that highly comparable pre-selection libraries diverge into bait-specific interaction profiles following Y2H selection.

To stringently eliminate nonspecific interacting fragments as well as those belonging to proteins that would not come into contact with Rab proteins, we expanded the set of protein fragments to filter out of the candidate list. This fragment-level analysis identified key fragments that came up in both vector-alone and some Rab datasets, indicating a nonspecific interaction, while a different fragment from that same protein was found to be strictly Rab-specific. These cases included nine candidates: PPP1R15A, RPN1, N4BP2L2, CREB3L4, PAF1, THOC5, PIAS2, KRT33B, and BANK1, each of which contributed a nonspecific background fragment as well as a distinct, bait-specific fragment scored as a genuine interaction (e.g., PPP1R15A; amino acids (aa) 64-165 were recurrently vector background, while PPP1R15A: aa 490-571 were uniquely and specifically recovered with Rab23).

The filtered dataset comprises 522 Rab prey interactions spanning 334 unique prey genes and all 36 Rab GTPases (**Supplementary Table 3**). Of these, 256 prey were recovered with a single Rab; 177 hits were GTP specific and 141 were both Rab and GTP specific (**Table 1**). The remaining 78 prey were recovered with two or more Rabs (**Table 2**). This scale of recovery, combined with the reproducibility and fragment level stringency of our filtering, demonstrates that DEEPN reliably resolves genuine, bait specific interactors from a large prey library.

Twenty-six of these interactors are already documented in BioGRID. Three, SNX13 and Rab9, SNX29 and Rab14, and MICAL-L1 and Rab15, had prior evidence only from proximity based methods (Go et al., 2021; Huttlin et al., 2021; Gonzalez-Lozano et al., 2025; Gillingham et al., 2019; Gaudreault et al., 2025; Antón-Plágaro et al., 2025). Our DEEPN Y2H data provide evidence for these interactions being direct. The 522 interactions reported here therefore represent a large, high confidence resource of likely Rab specific binding partners. The majority of these interactions have not been previously described and may serve unrecognized biological functions.

### Detection of Rab binding domains with multiple partners

DEEPN identified MICAL-L1 as an interactor for multiple Rab GTPases (**Figure 2A**), which showed a clear fragment enriched on Rab8, Rab10, Rab12, Rab13, Rab15 and Rab36 (**Figure 2B**). This fragment is defined by the peak of read-depth shown as well as discrete 5’ junction reads that span fusion point between the GBD vector and the prey plasmid insert. Three different fragments were inferred from DEEPN sequence data and used in binary Y2H assays to confirm interactions of these Rab GTPases (**Figure 2B**) and one of the strongest interacting fragments with the full panel of Rab GTPases (**Supplementary Figure 2A**). All of the Rab proteins identified by DEEPN as interacting in a nucleotide dependent manner were found to interact similarly in binary Y2H assays. Multiple Rab proteins bind MICAL-L1, mapping to a C-terminal region containing its trihelical bundle bivalent Mical/EHBP Rab Binding domain (bMERB). In some MICAL orthologs this domain can bind multiple Rabs (Rai et al., 2016; Esposito et al., 2018), with most homologs binding the Rab8 related subfamily (Kail et al., 2008). This family of proteins also contains a Calponin Homology (CH) domain and/or a LIM domain that can bind the bMERB domain to keep it in a closed, autoinhibited state, released when Rab competes for the CH/LIM binding site (Rai et al., 2024; Lin et al., 2024; Miyake et al., 2019). This sets up an activation mechanism in which Rab binding releases the actin binding CH domain, together with other N-terminal functional domains, to effect membrane trafficking and cell migration (Rahajeng et al., 2010; Sakane et al., 2018). MICAL-L1 is well-characterized for binding multiple Rab GTPases, making it a useful benchmark for evaluating DEEPN sensitivity and specificity. Five of 6 Rab proteins previously shown to interact with MICAL-L1 (Rab8, Rab10, Rab13, Rab15, Rab35, and Rab36) were recovered in DEEPN screens (**Figure 2B**). Recovery of MICAL-L1 across multiple GTP-locked Rab baits confirms the sensitivity and specificity of this approach.

**Figure 2.**
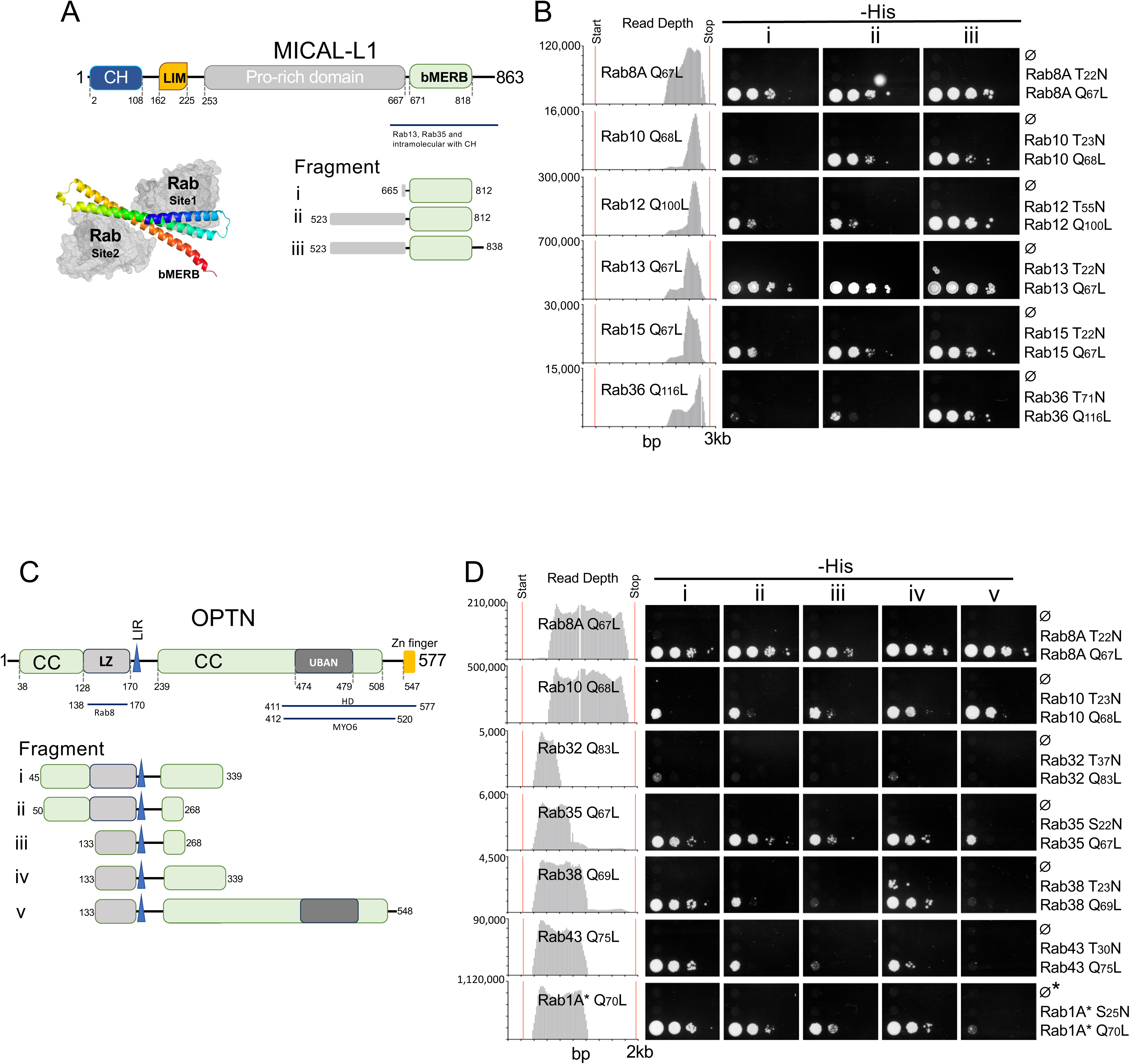
Rab interactions of MICAL-L1 and OPTN. **A.** Domain organization of MICAL-L1, showing the calponin homology (CH), LIM, proline-rich, and bivalent Mical/EHBP Rab Binding (bMERB) domains. The bMERB domain contains two Rab binding sites (Site 1 and Site 2) capable of simultaneous Rab engagement, as shown in the structural model. Model is AlphaFold structure of the MICAL-L1 bMERB domain with the 2 Rab10-binding sites of the MICAL1 bMERB domain superimposed. Three C-terminal fragments tested in binary assays are indicated (i: 665–812; ii: 523–812; iii: 523–838). **B.** DEEPN read depth profiles (left) showing enrichment of a bMERB-containing fragment under selective (−His) conditions for GTP-locked Rab8, Rab10, Rab12, Rab13, Rab15, and Rab36 baits. Binary Y2H spot assays (right) for fragments i, ii, and iii confirm GTP-dependent interactions. Serial dilutions were spotted onto permissive (+His) and selective (−His) plates; Ø denotes vector-only control. **C.** Domain organization of OPTN, showing two N-terminal coiled-coil (CC) regions flanking a leucine zipper (LZ), followed by a ubiquitin-binding domain (UBAN) and C-terminal zinc finger. Five overlapping fragments (i–v) spanning the N-terminal domain are indicated, with their boundaries shown relative to the domain map. **D.** DEEPN read depth profiles and binary Y2H spot assays for OPTN fragments i–v against GTP-locked Rab baits. Fragments containing aa 133-268 (encompassing the LZ) interact with Rab8, Rab10, and Rab35, consistent with the previously defined Rab8 binding site. Rab38 and Rab43 requires the N-terminal CC region C-terminal to the LZ (approximately aa 239-339) present in fragments i and iv. Rab1 interacts with fragments i-iv, but not v (aa133-548). Vector-only control is denoted (Ø).

DEEPN screens also identified Rab12, which prompted screening of the C-terminal bMERB region against the full Rab panel in both GTP and GDP-locked forms. Binary Y2H assays confirmed the interactions identified by DEEPN (**Supplementary Figure 2A**). Additional weaker interactions were also detected in binary assays (Rabs 9, 26, 32, 34, 35, and 38) but not in DEEPN. These had slow growth under His selective conditions, consistent with enrichment below the DEEPN threshold. A subset of Rabs was tested against different C-terminal fragments of MICAL-L1, all containing the three helix bMERB domain. Larger fragments that included unstructured sequence N-terminal to the bMERB domain interacted robustly with Rabs 8, 10, 12, 13, 15, and 36, but interaction was diminished for Rab10, 12, and 36 when that N-terminal region was removed. This is curious behavior since all fragments had the bMERB domain, but presented it in different contexts. Given the systematic way the Rab bait protein fusions were built, these data suggest the differential interactions across these baits **are** due to different binding modes on the bMERB domain. Indeed, previous structural analysis on a different bMERB domain identified 2 Rab binding sites (Rai et al., 2016) (**Figure 2A**). This would suggest Rab8 and Rab13 engage the bMERB domain in a way distinct from Rab10 and Rab36.

Optineurin (OPTN) is a 577-amino acid multidomain adaptor with a domain architecture comprising two N-terminal coiled-coil regions flanking a leucine zipper (LZ), followed by a LIM motif, additional coiled-coil domains, a ubiquitin-binding domain (UBD), and a C-terminal zinc finger (ZF) (**Figure 2C**). Its interaction with Rab8 is well established and structurally characterized: crystal structures of the OPTN LZ (aa 133-170) in complex with GTP-locked Rab8 define the core binding interface (Zhang et al., 2024), with deletion mapping placing the interaction within aa 141-209 (Sahlender et al., 2005; Ying and Yue, 2012). An interaction with Rab1 attributed to the C-terminal zinc finger domain has been reported in autophagosomal maturation (Song et al., 2018), and the glaucoma-associated M98K mutation has been shown to engage Rab12 in a pathway leading to autophagic retinal ganglion cell death (Sirohi et al., 2013), though the Rab12-binding site on OPTN has not been mapped. DEEPN screening identified OPTN enrichment on GTP-locked forms of multiple Rab GTPases (**Figure 2D**), with read-depth data indicating a different set of interacting fragments for Rab8 and Rab10, vs the other Rabs. Five overlapping fragments spanning amino acids (aa) (i: 45–339; ii: 50–268; iii: 133–268; iv: 133–339; and v: 133–548), were tested in binary Y2H assays against this set. Additionally, one fragment (aa 45-339) was tested against the whole Rab panel in binary Y2H assays (**Supplementary Figure 2B**). Rab8, Rab10, and Rab35 interacted with all five fragments, consistent with a binding determinant contained within the minimal shared region (aa 133-268), which encompasses the LZ and is consistent with the crystallographically defined Rab8 binding site. In contrast, Rab38 and Rab43 interacted robustly with fragment i (aa 45-339), showed reduced interaction with fragment ii (aa 50-268), and failed to interact with fragments iii (133–268) or v (133–548). This pattern implicates the N-terminal CC region C-terminal to the LZ (approximately aa 239-339) as an important determinant for Rab38 and Rab43 binding. Rab1 interacted with fragments i, ii, iii, and iv, and very weakly with fragment v (aa 133-548), despite v containing all sequences present in the binding-competent constructs. This raises the possibility that sequence in the 339–548 window (encompassing the UBD) may alter the conformation or accessibility of the Rab1-binding interface; notably, these N-terminal fragments identify a potential Rab1-interacting surface that is distinct from the C-terminal zinc finger implicated by prior deletion analysis (Song et al., 2018), suggesting the interaction may be more complex than previously appreciated. Rab3 showed weak interaction limited to fragment i, and Rab32 had only very weak interaction with i and iv. The interactions with Rab10, Rab35, Rab38, Rab43, and Rab3 have not been previously described. Together, these data establish OPTN as a multi-Rab effector with at least two Rab-binding surfaces within its N-terminal domain architecture, the structural basis and functional significance of which remain to be determined.

RABEP1 (Rabaptin-5) is a scaffold organized into four coiled-coil segments (CC1-1, CC1-2, CC2-1, CC2-2) that connects the Vps9-domain GEF RABGEF1 (Rabex-5) to Rab4-containing compartments through Rab4 and Rabex-5 binding domains housed in its N-terminal and C-terminal regions (**Figure 3A**). (Stenmark et al., 1995; Horiuchi et al., 1997). RABEP1 also binds GGA adaptors through a region spanning approximately aa 301-449, linking early endosomal sorting to trans-Golgi network retrieval (Mattera et al., 2003). Prior two-hybrid and biochemical studies have mapped Rab4-GTP binding to two distinct regions in the N-terminal portion of RABEP1: aa 5-135 (Vitale et al., 1998) and aa 140-295, with aa 187-226 necessary for binding (Deneka et al., 2003; Korobko et al., 2005). The adjacent CC1-2 helix (aa 216-318) binds Rab5-GTP (Korobko et al., 2006). DEEPN also identified these distinct Rab binding regions, which were confirmed in binary assays (**Figure 3B**). Rab4-GTP enriched a RABEP1 fragment encompassing aa 62-261, whereas Rab5-GTP enriched a fragment encompassing aa 222-393 (**Figure 3B**). The Rab4-binding fragment spans both previously mapped Rab4 binding regions, and since aa 222-393 do not bind Rab4, the contact site lies within aa 62-221; the data do not resolve between the two sites. Rab5 binding resolves to aa 222-318 (**Figure 3B**).

**Figure 3.**
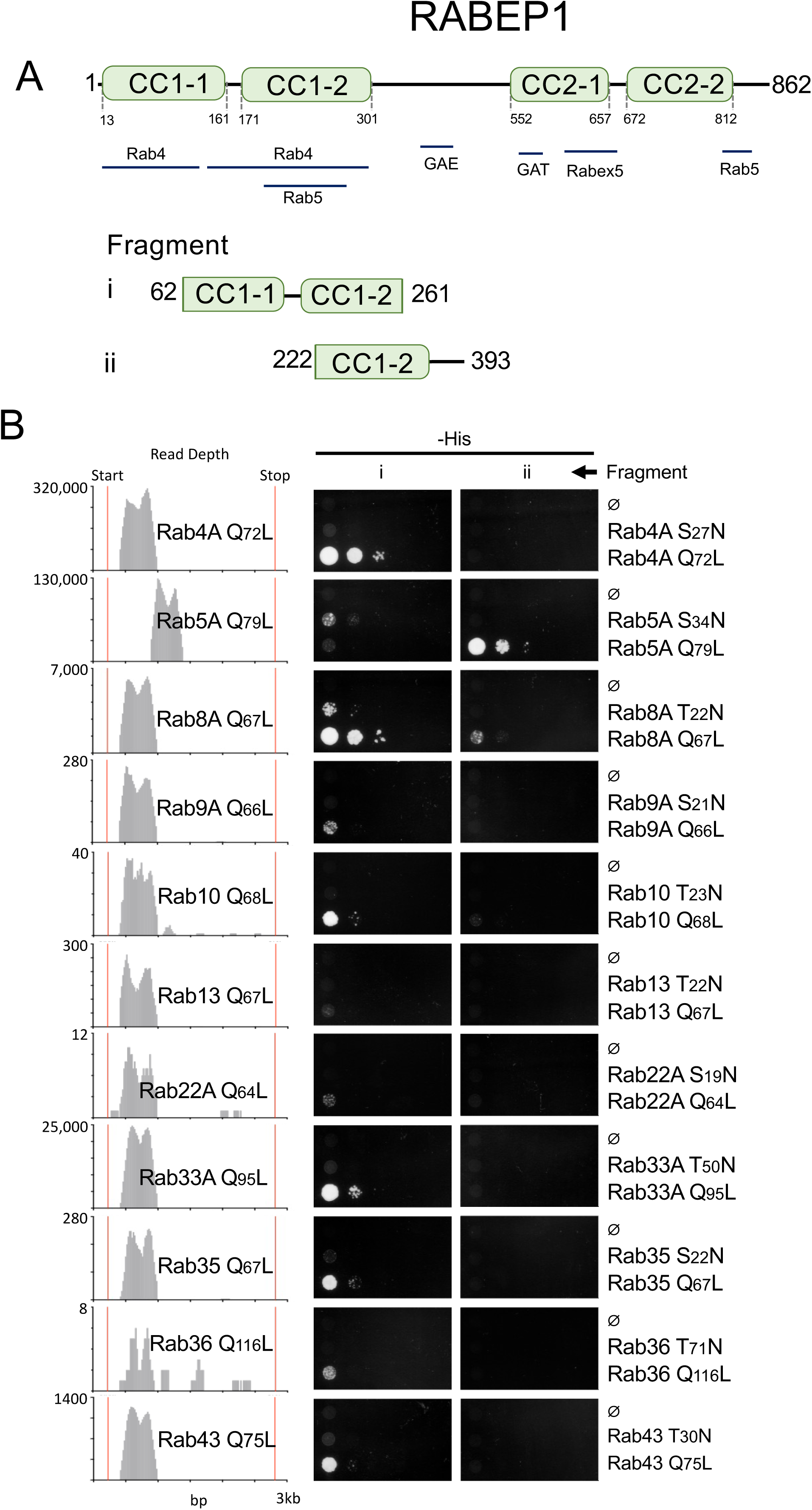
Interactions of RABEP1. **A.** Domain organization of RABEP1 (Rabaptin-5), showing four coiled-coil segments (CC1-1, CC1-2, CC2-1, CC2-2) and previously mapped binding sites for Rab4, Rab5, GGA adaptors, Rabex-5 (RABGEF1), and the C-terminal Rab5-binding domain (R5BD). Two DEEPN fragments are indicated: fragment i (aa 62-261, spanning CC1-1 and CC1-2) and fragment ii (aa 222-393, within CC1-2). Comparison of DEEPN fragment boundaries with previously mapped binding regions. **B.** DEEPN read depth profiles (left) and binary Y2H spot assays (right) for fragments i and ii against a panel of Rab baits. Fragment i is enriched by Rab4, Rab8, Rab9, Rab10, Rab13, Rab22, Rab33, Rab35, Rab36, and Rab43, all in their GTP-locked conformation. It also is weakly enriched by Rab5 in its GDP-locked conformation. Fragment ii shows strong interaction with Rab5 and weak interaction with Rab8 in binary assays. Ø denotes vector-only control.

DEEPN also identified that the Rab4-binding region could interact with several other Rabs, all in their GTP-bound conformation (Rab8, Rab10, Rab13, Rab33, Rab35, and Rab43). This was confirmed with the full panel of Rab GTPases in Y2H binary assays (**Supplementary Figure 3**), which also uncovered weak interactions with Rab9, Rab22, and Rab36 that did not meet the identification thresholds we set for DEEPN, but which clearly showed selection of the interacting fragment by read depth profiles. This expanded repertoire of interacting Rabs is consistent with proximity proteomics assays that place these Rabs near where RABEP1 functions (Go et al., 2021; Liu et al., 2018; Wilson et al., 2023). The panel of Rab binary interactions with the Rab5 binding fragment (aa 222-393) confirmed strong interaction with Rab5-GTP, and weak interaction with Rab8-GTP. Interaction of Rab8 with this more distal site was not detected by DEEPN, most likely because competition by the more proximal CC1-1 Rab binding site inhibited enrichment.

One model for how RABEP1 functions as a feed-forward mechanism that allows activated Rab4 to recruit RABEP1 to membranes, to initiate the activation of Rab5 via interaction with Rabex-5, which causes accumulation of Rab5-GTP that stabilizes RABEP1 on the membrane through its Rab5 binding site adjacent to the Rab4 binding site (Kälin et al., 2015; Kälin et al., 2016). One implication from our data is that other Rabs besides Rab4 can localize Rab5 exchange activity across a broader range of compartments throughout the endocytic pathway. Our data also showed the adjacent Rab5-GTP binding domain on the CC1-2 helix is remarkably specific for Rab5, which implies that Rab5 placement there, versus a broader array of Rab proteins, might serve a more specific function than to simply stabilize RABEP1 to Rab5 compartments. One possibility is that Rab5 occupancy adjacent to the more promiscuous CC1-1 Rab binding site(s) competes for Rab4 (and other) binding, working as a shut-off mechanism to dislodge RABEP1 from these other Rab-containing membranes.

### Additional Interactions

Some candidates when tested in binary screens across the full Rab panel, found a wider range of interactions that were missed by DEEPN, likely because they were outcompeted in that particular population cohort. This underscores that relative enrichment between bait groups vs just achieving a threshold would be problematic, and reinforces the types of statistical analysis performed here where a hit is defined by a combination of threshold gates. For instance, DEEPN found APPL2 interacts with Rab31 and Rab22 (**Supplementary Figure 4**). Binary Y2H assays confirmed this, but also identified Rab5, all three of which have been previously reported interactors of APPL2 (King et al., 2012). DEEPN identified Rab11FIP2 as an interactor for both Rab11 and Rab14 (**Supplementary Figure 5**). Binary Y2H assays confirmed this, but also extended the interaction profile to include Rab22, Rab31, and Rab43. DEEPN data for Rab31 showed a weak signal but fell below significance (p > 0.1), whereas Rab22 and Rab43 were well below any threshold for enrichment filtering. DEEPN found RUFY2 interacted with Rab38-GTP and Rab33-GTP, but missed Rab5-GDP and Rab33-GDP, both of which fell below the threshold for percentage of in-frame junctions (**Supplementary Figure 6**). DEEPN identified IFT81 as an interactor for Rab5-GTP and Rab31-GTP, but missed a Rab22-GTP interaction that was observed by binary Y2H assays. DEEPN enrichment data gave no indication that Rab22-GTP was close by any filter criteria (**Supplementary Figure 7**). By contrast, DEEPN agreed with binary Y2H panels on the exclusive Rab-specific interactions of SNX29 with Rab14-GTP, HPS4 with Rab9-GTP, and DAB2IP with Rab36-GTP (**Supplementary Figures 8, 9, and 10**).

We also found BBS7 and SYN3 were both identified as Rab interactors by DEEPN, while not enriching on any p*TEF1*-GBD vector alone dataset. However, they were eliminated by an additional layer of stringency of also excluding candidates that enriched on a slightly different Gal4-BD bait plasmid, pGBKT7, distributed as part of the Clontech/Takara Y2H system. Vector-only data from pGBKT7 were previously described (Wiemann et al., 2016). This plasmid is a Gal4-DNA binding domain fusion is a high copy plasmid with different linker than characteristics than the p*TEF1*-GBD low-copy plasmid used throughout this study. Binary Y2H assays indicate that both proteins do have specific Rab interactions, and for BBS7, some binary assays also showed a low level of vector-alone growth, consistent with the vector-associated background DEEPN detected (**Supplementary Figure 11**). Together, this suggests that treating any enrichment on either vector background as disqualifying (e.g., the p*TEF1*-GBD vector developed here or the pGBKT7 vector described elsewhere) sets a high bar that may exclude solid interactors along with genuine background. For instance, DEEPN found BBS7 (aa 391-486) interacted with Rab8 (GTP and GDP), Rab12-GTP, Rab13-GTP, and Rab19-GTP, which were confirmed in binary Y2H assays. DEEPN also found an interaction of SYN3 (aa 61-468) with Rab26-GTP, which was verified by binary Y2H assays (**Supplementary Figure 12**), but it was eliminated from the filtered list of interactors due to enrichment on pGBKT7 vector-alone samples. These data indicate that the stringency applied here is calibrated to minimize false positives, accepting some false negatives as a consequence.

### SNX RGS interaction

SNX13 and SNX14 belong to a subfamily of sorting nexins that have an RGS (regulator of G protein signaling) domain, known canonically to accelerate GTP hydrolysis by G-alpha GTPases (Amatya et al., 2021; Hollinger and Hepler, 2002; Masuho et al., 2020). Both carry N-terminal transmembrane domains that for SNX14 localize it in the ER, and a PI3P-binding PX domain thought to help it tether to endosomes (Hariri et al., 2019; Lu et al., 2022). Both have been implicated in lipid droplet biogenesis (Hariri et al., 2019; Ugrankar et al., 2020; Lu et al., 2022), and loss-of-function mutations in SNX14 cause SCAR20, an autosomal recessive spinocerebellar ataxia (Zhou et al., 2024; Ugrankar et al., 2018). We found that a fragment containing the RGS domain of SNX13 was enriched prey on multiple Rab proteins in DEEPN screens, including Rab5 and Rab9 with Rab9 interacting in both its GDP and GTP conformation (**Figure 4A**). This was confirmed in binary Y2H interactions against this set of Rab GTPases (**Figure 4B**) as well as across the complete panel of Rab proteins (**Supplementary Figure 13A**). SNX13 was originally characterized as a GAP for Gαs (Zheng et al., 2001), however, robust GAP activity for Gαs has not been tied to SNX13’s cellular function. RGS domains classically bind Galpha GTPases mostly through the Gα G-domain, which is shared across small molecular weight GTPases such as the Rab family. These structural parallels implied that these SNX RGS domains may serve more generally as Rab interactors (**Figure 4E**). To examine this, we tested the RGS domain of SNX14 in binary Y2H screens (**Figure 4C** and **Supplementary Figure 13B**), which showed the SNX14 RGS domain interacted with Rab11, Rab32, Rab9 and Rab38 in their GTP-bound conformation, but not with Rab5 in contrast to the SNX13 RGS domain. We also tested whether an alternate GTP-locking substitution behaved equivalently to Q70L. Here we used a Rab11 S20V, which is also locked into a GTP-bound conformation (Choi et al., 2016). Unexpectedly, we found that Rab11 S20V did not interact with SNX14-RGS domain (**Supplementary Figure 13C**). However, this appeared to be a specific failure of the residue change itself as S20 lies near the SNX14 binding interface in the structural model we generated (**Figure 4E**). Indeed, introducing S20V into the Q70L background largely eliminated the otherwise strong Q70L interaction (**Supplementary Figure 13C**).

**Figure 4.**
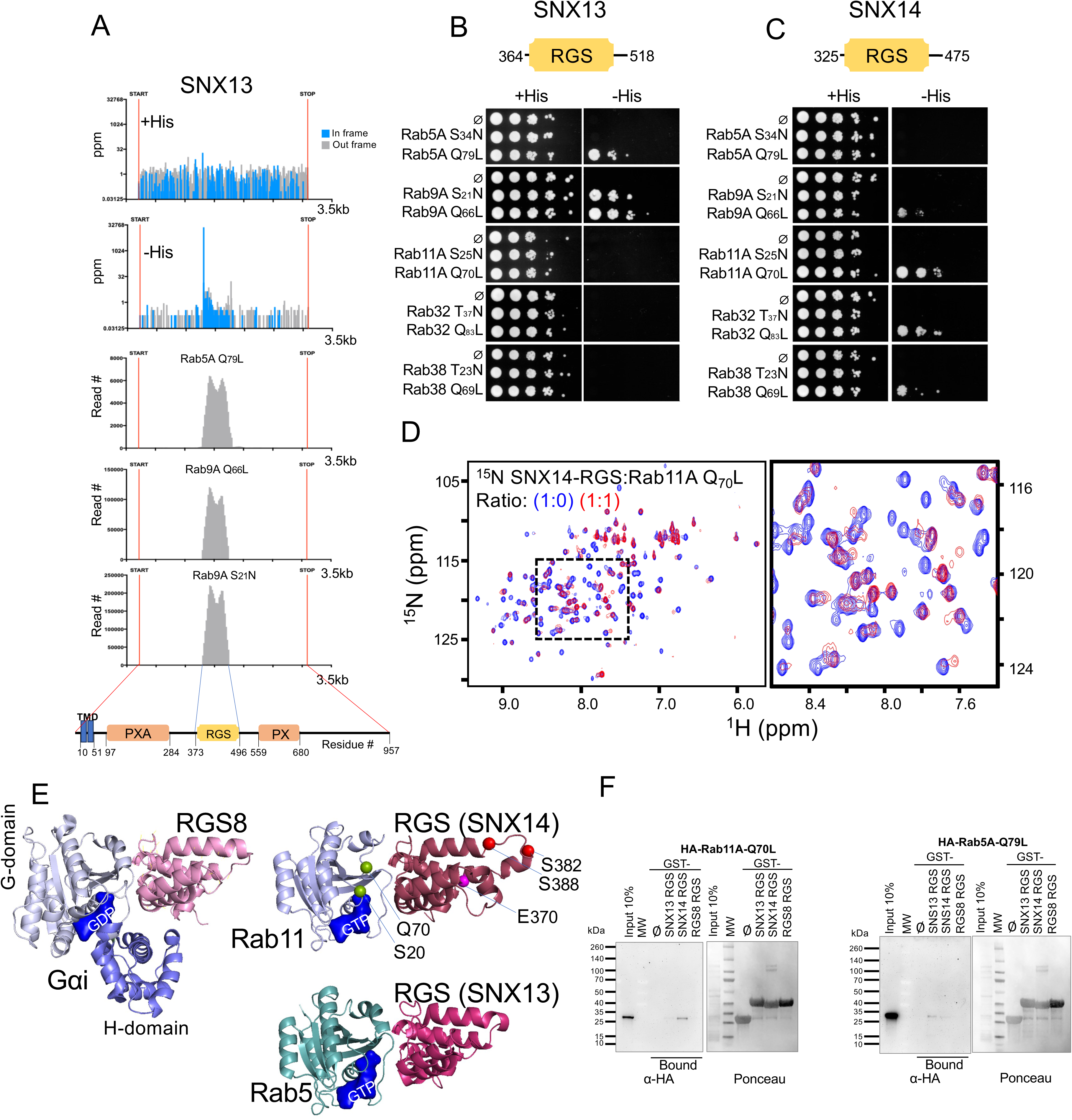
Rab interaction with the RGS domains of SNX13 and SNX14. **A.** DEEPN read depth profiles for SNX13 showing enrichment of an RGS domain-containing fragment under selective (−His) conditions with Rab5 Q79L, Rab9 Q66L, and Rab9 S21N baits, and the corresponding non-selective (+His) profile. Domain organization of SNX13 is shown below, including the transmembrane (TMD), PXA, RGS, and PX domains. **B.** Binary Y2H spot assays for the SNX13 RGS domain (aa 364-518) against GTP-locked and GDP-locked Rab5, Rab9, Rab11, Rab32, and Rab38. Rab5 binding is GTP-dependent; Rab9 binding occurs with both nucleotide states. Vector-only control is denoted (Ø). **C.** Binary Y2H spot assays for the SNX14 RGS domain against the same Rab panel. SNX14 RGS binds Rab9, Rab11, Rab32, and Rab38 in a GTP-dependent manner. **D.** 2D ¹H-¹⁵N HSQC spectra of ¹⁵N-labeled SNX14 RGS alone and with unlabeled Rab11 Q70L at 1:0 and 1:1 molar ratios. Chemical shift perturbations and resonance broadening (boxed region shown at right, expanded) confirm direct physical interaction. Vector-only control is denoted (Ø). **E.** Structural comparison of RGS–GTPase complexes. Left: crystal structure of RGS8 bound to activated Gαi3 (PDB: 2ODE), illustrating the canonical RGS–Gα interface in which the RGS domain engages switch regions of the Gα GTPase (G) domain. Right: AlphaFold2 models of the SNX14 RGS domain bound to Rab11-GTP (top) and the SNX13 RGS domain bound to Rab5-GTP (bottom), with key aa E370, Q70, S20, S382, and S388 indicated. Though Rab GTPases lack the α-helical (H) domain of Gα, RGS domains engage the G domain, which is conserved across small GTPases. **F.** GST pulldown assays. HA-tagged Rab11 Q70L (left) or HA-tagged Rab5 Q79L (right) was incubated with GST alone (Ø), GST-SNX13 RGS, GST-SNX14 RGS, or GST-RGS8 immobilized on glutathione beads. Bound material was detected by anti-HA immunoblot; Ponceau staining confirms equivalent GST loading.

Direct binding of the RGS domains to GTP-bound Rab proteins was confirmed by GST pulldown. GST-SNX14 RGS bound recombinant Rab11 Q70L, whereas no binding was observed for the RGS domain of RGS8 fused to GST, and little binding of Rab11 by GST-SNX13-RGS. In contrast, GST-SNX13-RGS bound Rab5 in its GTP conformation (Q79L) with less binding observed for SNX14-RGS and little binding by RGS8 (**Figure 4F**). Binding was further examined using NMR spectroscopy. The 2D ¹H-¹⁵N HSQC spectra of ¹⁵N-labeled SNX14 RGS showed both chemical shift perturbations and selective peak broadening upon addition of unlabeled Rab11 Q70L, independently confirming a direct physical interaction (**Figure 4D**). The quality of these proteins is shown in **Supplementary Figure 13F**.

Given the architecture, shared Rab-binding properties, and previously reported ER localization, we sought to characterize SNX13 and SNX14 further using GFP and mCherry fusion proteins (**Figure 5A**). Co-immunoprecipitation experiments showed that each protein self-associates and that SNX13 and SNX14 associate with one another (**Figure 5B**). Though these experiments cannot definitively distinguish dimeric from higher-order complexes, they do imply that both proteins form oligomers beyond simple dimers. SNX13 and SNX14 co-localized extensively by fluorescence microscopy, consistent with the co-immunoprecipitation data (**Figure 5C**). And each protein partially overlapped with the ER marker GFP-KDEL without fully coinciding with it, consistent with association with an ER subdomain rather than bulk ER (**Figure 5C**) and consistent with previous localization studies (Hariri et al., 2019; Lu et al., 2022).

**Figure 5.**
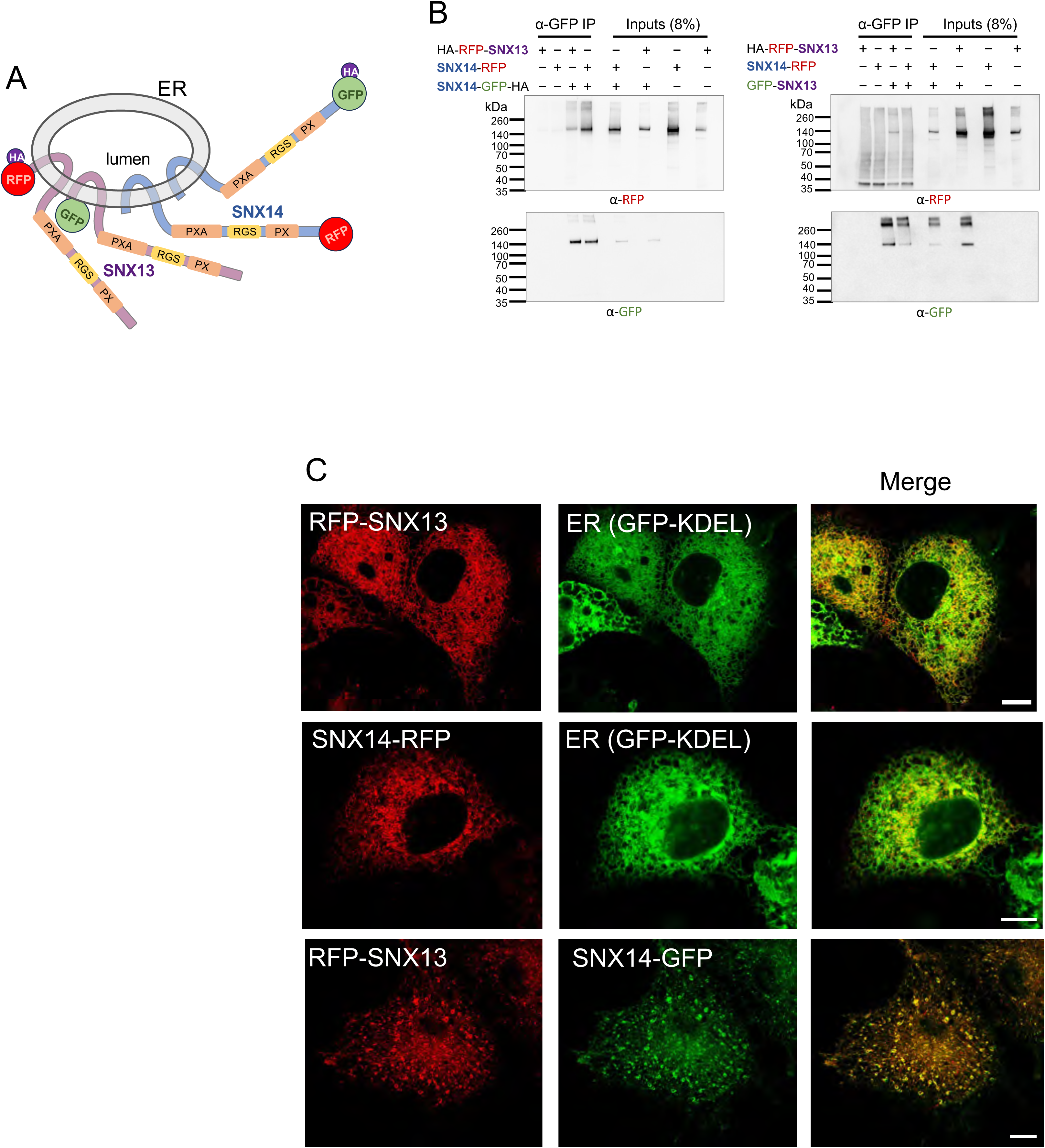
Oligomerization and co-localization of SNX13 and SNX14. **A.** Schematic of SNX13 and SNX14 domain topology at the ER membrane, showing cytoplasmic orientation of the RGS, PXA, and PX domains and the intralumenal loop. Tagged constructs used for interaction and localization studies are indicated. **B.** Co-immunoprecipitation of SNX13 and SNX14 from COS-7 cells. The indicated combinations of HA-RFP-SNX13, SNX14-RFP, and SNX14-GFP-HA were co-expressed; GFP nanobeads were used for immunoprecipitation and blots were probed for RFP and GFP. SNX14 homo-dimerizes and co-precipitates SNX13. Right panels: equivalent experiment showing SNX13-GFP co-precipitating SNX14-RFP. **C.** Confocal fluorescence microscopy of COS-7 cells. RFP-SNX13 and SNX14-RFP each partially co-localize with the ER marker GFP-KDEL, consistent with association with an ER subdomain. RFP-SNX13 and SNX14-GFP co-localize extensively, consistent with shared complex formation at the ER. Scale bar=10 μm.

We also found evidence that Rab interaction might be regulated by phosphorylation. Previous studies showed that SNX14 is phosphorylated at S382 and S388 by PKA in response to serotonin signaling (Ha et al., 2015). Phosphomimetic substitutions (S382D and S388D) at these positions selectively disrupted Rab32 binding in binary Y2H assays. Yet Rab11 and Rab9 interactions were unaffected (**Figure 6A**) and protein levels of these mutants were comparable to WT RGS (**Supplementary Figure 4D**). These data implicate phosphorylation as a mechanism for modulating specific Rab partner interactions of the SNX RGS domain.

**Figure 6.**
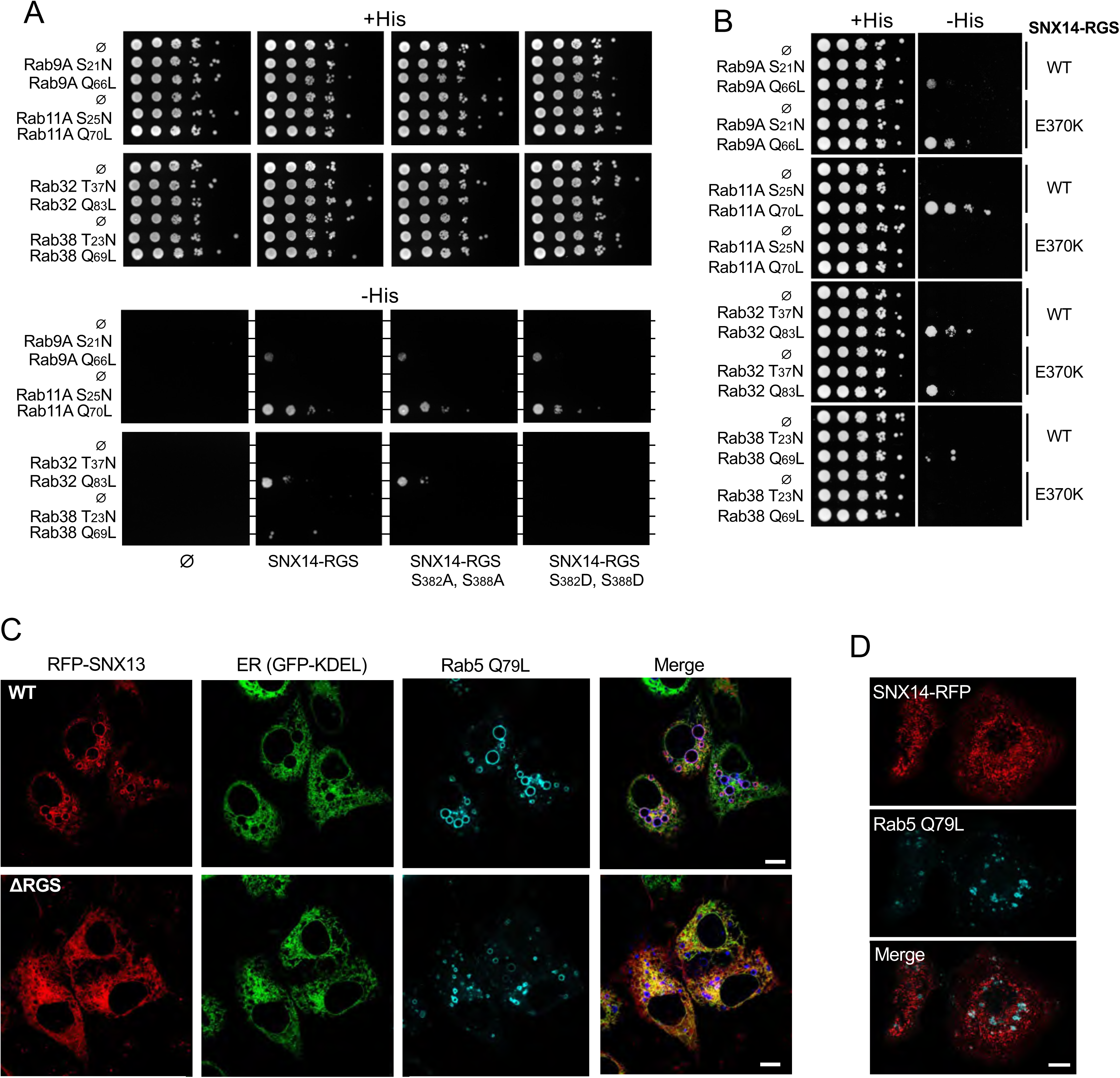
Functional link of SNX14 to Rab GTPases. **A.** Binary Y2H spot assays comparing WT SNX14 RGS, phospho-null (S382A, S388A), and phosphomimetic (S382D, S388D) variants against GTP- and GDP-locked Rab9, Rab11, Rab32, Rab38, and vector alone (Ø). Phosphomimetic substitution selectively disrupts Rab32 binding while Rab11 and Rab9 interactions are unaffected, identifying phosphorylation as an independent mechanism modulating Rab partner selectivity through the SNX14 RGS domain. **B.** Binary Y2H comparing WT SNX14 RGS and E370K against Rab9, Rab11, Rab32, and Rab38 in GTP-locked and GDP-locked forms. E370K abolishes Rab11 binding, leaves Rab32 largely unchanged, and modestly increases Rab9 interaction. Vector-only control is denoted (Ø). **C.** Confocal fluorescence microscopy of COS-7 cells expressing full-length RFP-SNX13 (WT or ΔRGS) together with GFP-KDEL and BFP-Rab5 Q79L. WT SNX13 localizes prominently to enlarged Rab5 Q79L-positive endosomes; ΔRGS SNX13 fails to accumulate on these structures, demonstrating that RGS-mediated Rab5 binding is required for Rab5 endosome recruitment. Scale bar=10 μm. **D.** SNX14-RFP co-expressed with BFP Rab5 Q79L showing it does not localize to exaggerated endosomes. Scale bar=10 μm.

The disease-causing missense variant E370K of SNX14 has previously been identified in SCAR20 patients (Sait et al., 2022). E370 maps to the RGS domain (**Figure 4E**) and we found that in binary Y2H assays, the E370K substitution abolished binding to Rab11, left binding to Rab32 largely unchanged, and modestly increased binding to Rab9 (**Figure 6B, Supplementary Figure 13C**). A single substitution producing three distinct outcomes across three Rab partners argues against simple destabilization of the RGS fold and instead identifies E370 as a residue that contributes differentially to each binding interface, consistent with its position near the predicted Rab11 contact surface in the AlphaFold model (**Figure 4E**). Protein levels of the E370K prey fusion were reduced relative to wild type (**Supplementary Figure 13D**), consistent with some instability, though the remaining protein supported robust Rab9 interaction.

Previous experiments partially localized SNX13 to endosomes (Zheng et al., 2001; Gullapalli et al., 2006). To determine whether Rab–RGS binding was relevant to its cellular function and localization we examined the contribution of the RGS domain. Full-length SNX13 localized prominently to enlarged endosomes induced by expression of BFP-Rab5 Q79L, in some cases with adjacent ER membrane apparent near these structures (**Figure 6C**). However, SNX13 lacking its RGS domain failed to localize efficiently to these endosomes (**Figure 6C**), showing that RGS-mediated Rab binding helps recruit SNX13 to this compartment. In contrast, SNX14, whose RGS domain does not engage Rab5, did not accumulate on Rab5 Q79L endosomes (**Figure 6D**).

## DISCUSSION

We screened 36 human Rab GTPases against a fragment library built from Human ORFeome v8.1, identifying 522 interactions with 334 prey proteins across GTP- and GDP-locked conformations (Pashkova et al., 2016; Peterson et al., 2018; Peterson and Piper, 2021). This dataset is comprehensive because the same prey library was screened against every bait under identical growth and selection conditions. A newly made library of fragments derived from the human ORFeome allows survey of distinct domains that may be more free to interact outside the context of their parent protein. Both the reproducibility of the prey plasmid library population and the identification of the fragments sufficient for interaction are enabled by high-throughput sequence data that can be parsed with the updated software reported here. The updated DEEPN software runs on Apple Silicon, uses a DESeq2-based statistical model, and includes a FragFinder module for reconstructing and annotating interacting fragments. Broader accessibility to these methods is possible given that the code and rationale for it are available publicly and can be adapted with the help of AI models that have facility with coding and implementation. The depth of interactome exploration is still limited to the variety of prey library plasmid elements available. The new human ORFeome fragment library, housed in a streamlined low-copy plasmid, has more complexity than some of the commercially available Y2H cDNA libraries, however, the ORFeome v8.1 used here does not fully represent at least one ORF for all genes, and additional libraries using more recent ORFeomes could be helpful. Another limitation is that since fragments are ligated randomly, ∼1 in 6 random fragments is in productive reading frame. Supplementary approaches, such as fusion of a P2A-Ura3 cassette 3′ of the insert, could help select for a library that orients fragments in the proper reading frame.

Proteome-scale Y2H screens have indeed captured Rab interactions, however, these were not designed to capture nucleotide-specific interactions, which is a key aspect for identifying the biologically relevant Rab interactomes (Rual et al., 2005; Stelzl et al., 2005; Ewing et al., 2007). Proximity proteomics has provided Rab interactions in a biological context where their nucleotide states would operate (Antón-Plágaro et al., 2025; Gillingham and Munro, 2014; Gillingham et al., 2019; van Vliet et al., 2025; Gaudreault et al., 2025). And these approaches were complemented by cross-linking enabled proteomic analysis of purified endosomes that captures endosomal Rab interactions (Gonzalez-Lozano et al., 2025). Our DEEPN data capture interactions at fragment resolution with defined nucleotide states in a context that likely demands direct protein∼protein interaction. Thus, where there is overlap in the identification by DEEPN vs these additional approaches, DEEPN indicates direct interaction on a particular protein subdomain.

One measure of the robustness of DEEPN is its ability to recover true interactions of the same protein or protein domain across multiple Rabs. This was evident for several well-characterized Rab effectors. The bMERB domain of MICAL-L1 (Kail et al., 2008) interacted with Rab8, Rab10, Rab13, Rab15, and Rab36, consistent with its established role as a multi-Rab binding domain, while also identifying Rab12 as an additional partner. RABEP1, the founding Rab5 effector (Stenmark et al., 1995; Vitale et al., 1998), showed the expected interactions with Rab4 and Rab5, but its Rab4-binding region also interacted with Rab8, Rab10, Rab13, Rab33, Rab35, and Rab43. DEEPN also recovered the established interaction between OPTN and Rab8 (Hattula and Peränen, 2000; Zhang et al., 2024), while extending its Rab-binding profile to Rab1, Rab10, Rab32, Rab35, Rab38, and Rab43. These interactions were strongly GTP-selective, providing an additional internal measure of specificity. Table 2 shows additional candidate proteins that interact with more than one Rab. None of these interacting fragments were enriched in any of 17 independent vector-alone selections, adding confidence that they represent bona fide multi-Rab interactors. We found that the RGS domains of SNX13 and SNX14 differentially interact with Rab GTPases. These proteins belong to a class containing SNX-PXA-RGS-PXC domains that includes SNX25 (Amatya et al., 2021). Interaction of an RGS domain with a Rab GTPase does have one precedent, where the RGS domain of D-AKAP2 interacts with Rab4 and Rab11 (Eggers et al., 2009). RGS binding to Rabs reflects convergent use of a conserved interface: both Gα and Rab GTPases share the same G-domain fold. The concave helical bundle of the RGS fold contacts the Gα switch regions to impart GAP activity, and the equivalent switch I and switch II elements of Rab GTPases adopt a competent binding surface upon GTP loading. SNX25 also associates with Rab11, implying its RGS domain mediates Rab interaction as well (Maruzs et al., 2023). Whether RGS-domain-mediated Rab binding extends beyond SNX-RGS proteins is an open question. The 20 canonical mammalian RGS proteins have established Gα substrates (Masuho et al., 2020) but have not been systematically tested for Rab interactions. RGS-domain-containing proteins outside this canonical family, beyond the SNX-RGS members characterized here, are additional candidates.

SNX13 and SNX14 self-associate, forming oligomers that may combine their distinct properties (**Figure 5**). DEEPN identified GTP-selective Rab5 interactions through the RGS domain of SNX13, and consistent with this, SNX13 colocalizes with exaggerated Rab5-positive endosomes in a manner that depends on its RGS domain, as deletion of the RGS abolishes this recruitment (**Figure 6C**). This extends earlier colocalization of SNX13 with the Rab5 effector EEA1 (Gullapalli et al., 2006) and shows that RGS-mediated Rab binding contributes directly to endosome association. Together, PI3P recognition by the PX domain and GTP-Rab5 engagement by the RGS domain provide a dual-anchor mechanism tethering the ER to the endosomal surface. What such ER-endosome tethering accomplishes in this context is not fully resolved. SNX14 has been implicated in lipid droplet biogenesis (Hariri et al., 2019; Ugrankar et al., 2020), and ER-endosome contact sites have been proposed to support lipid transfer relevant to both multivesicular body sorting and lipid droplet formation (Hugenroth and Bohnert, 2019).

Whatever the molecular role of Rab-RGS domain might be, SNX14 plays an important role physiologically. Loss-of-function mutations cause SCAR20, a recessive spinocerebellar ataxia (Zhou et al., 2024; Ugrankar et al., 2018). Evidence that Rab binding is functionally important at the physiological level comes from a SCAR20-associated missense mutation, E370K, which falls within the Rab binding RGS domain region (Sait et al., 2022). We find that the E370K variant selectively loses its ability to bind Rab11 while retaining interaction with Rab9. Levels of the E370K prey fusion protein in yeast are reduced relative to wild type, consistent with some degree of instability. Yet these reduced levels are sufficient for robust Rab9 interaction, indicating that the loss of Rab11 binding cannot be attributed to simple misfolding. The selective loss of Rab11 binding by this disease variant implicates association with Rab11 recycling endosomes as a contributor to this ataxia.

Overall, this DEEPN dataset is striking because many of the interactions it found have not been previously reported. Thus, despite decades of work to determine the Rab interactome using a variety of high-throughput methods including recent proximity MS techniques, the full extent of Rab biology may well be still incomplete. Certainly, some of the interactions described here that pass through the stringent filters we applied may be biochemically relevant yet ultimately lack biological relevance. Yet others, such as the ones further explored here, are likely to indeed have definable Rab-specific roles. The dataset adds to an accumulating body of Rab interactome work that, taken together, moves toward filling in the mechanistic basis of the many cellular processes that depend on Rab proteins.

## Author contributions

TP and RP were responsible for generation of experimental and/or computational data. CP was responsible for NMR data and recombinant protein production. MS contributed to statistical analysis and programming. All were responsible for manuscript editing and data interpretation.

## Acknowledgements

We acknowledge the University of Iowa personnel and instrumentation in the IIHG Genomic Sequencing, the Carver College of Medicine NMR, and Protein & Crystallography core facilities, supported by the Roy J. and Lucille A. Carver College of Medicine and grants from the Roy J. Carver Charitable Trust. We thank Einat Snir for careful work on the Illumina sequencing of library amplicons and major contribution by Venkatramanan Krishnamani on how to update DEEPN software. This work was supported by NIH R01GM058202 to RCP. RCP were supported by the Roy J. Carver Charitable Trust.

---

**Supplementary Table 1. Plasmids. Plasmids used in this study**

**Supplementary Table 2. Non-specific, non-relevant interactors.** Compiled list of prey fragments eliminated from the list of potential interacting partners and the criteria used for their elimination.

**Supplementary Table 3. Candidate Rab interactors.** Interactors are shown for each Rab GTPase, designated for those that bind the GTP and/or the GDP conformation. Interactions previously described in BioGrid are in blue.

**Supplementary Figure 1.**
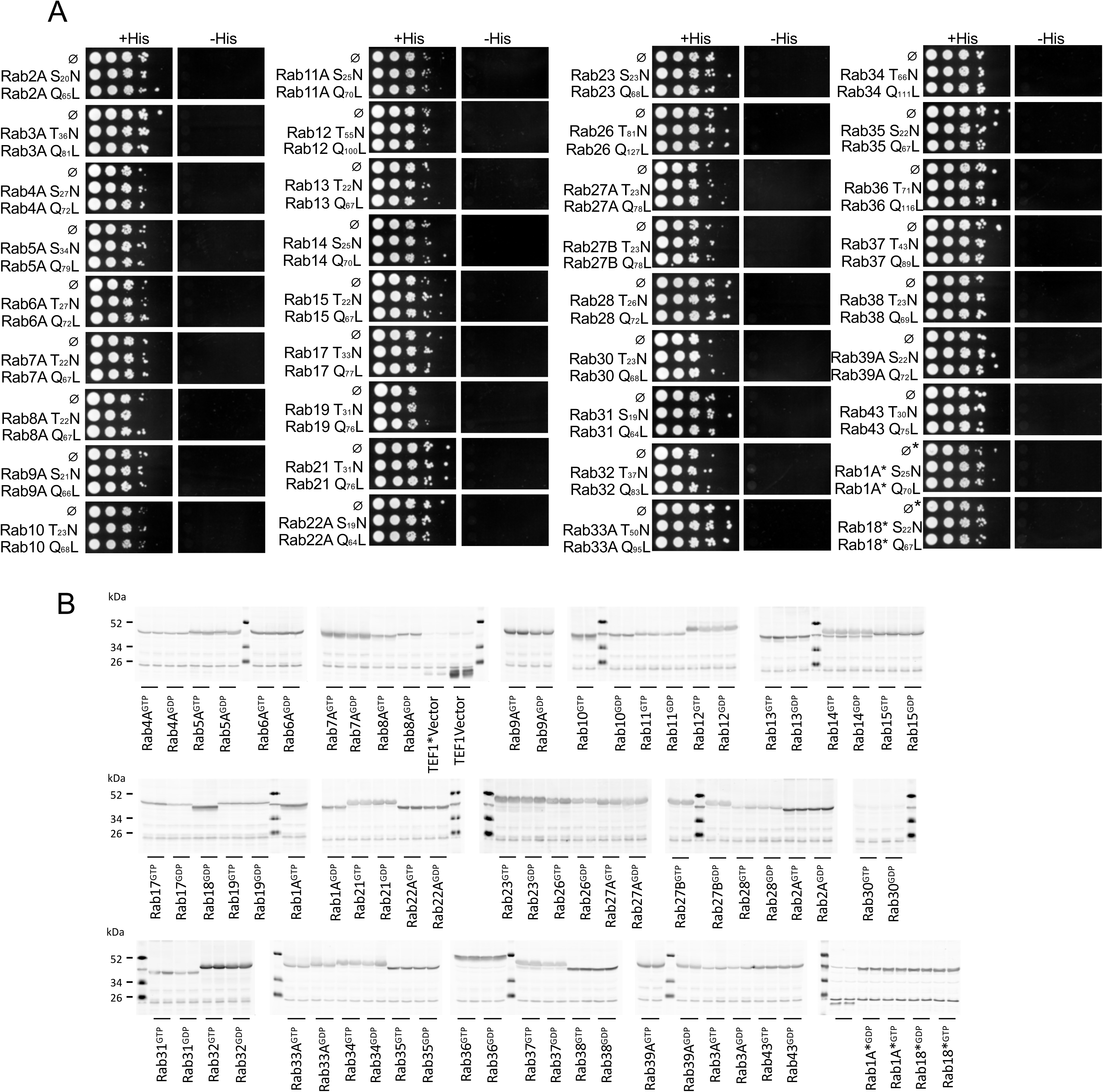
Absence of self-activation by Rab GTPase baits. **A.** Binary Y2H spot assays of each Rab GTPase bait (GTP-locked Q>L or GDP-locked S/T>N) paired with empty Gal4 activation domain vector, spotted onto permissive (+His) and selective (−His) plates. Absence of growth on −His confirms that no bait construct self-activates the reporter. Asterisks (*) denote Rab1 and Rab18, expressed from the reduced-strength *Tef1*\* promoter. Vector-only control is denoted (Ø). **B.** Anti-myc immunoblots of yeast transformants expressing the indicated p*TEF1*-GBD and p*Tef1*\* fusion proteins. Two colonies of each transformant pool were analyzed. Only transformants verified for expression of the bait fusion were used in DEEPN runs.

**Supplementary Figure 2.**
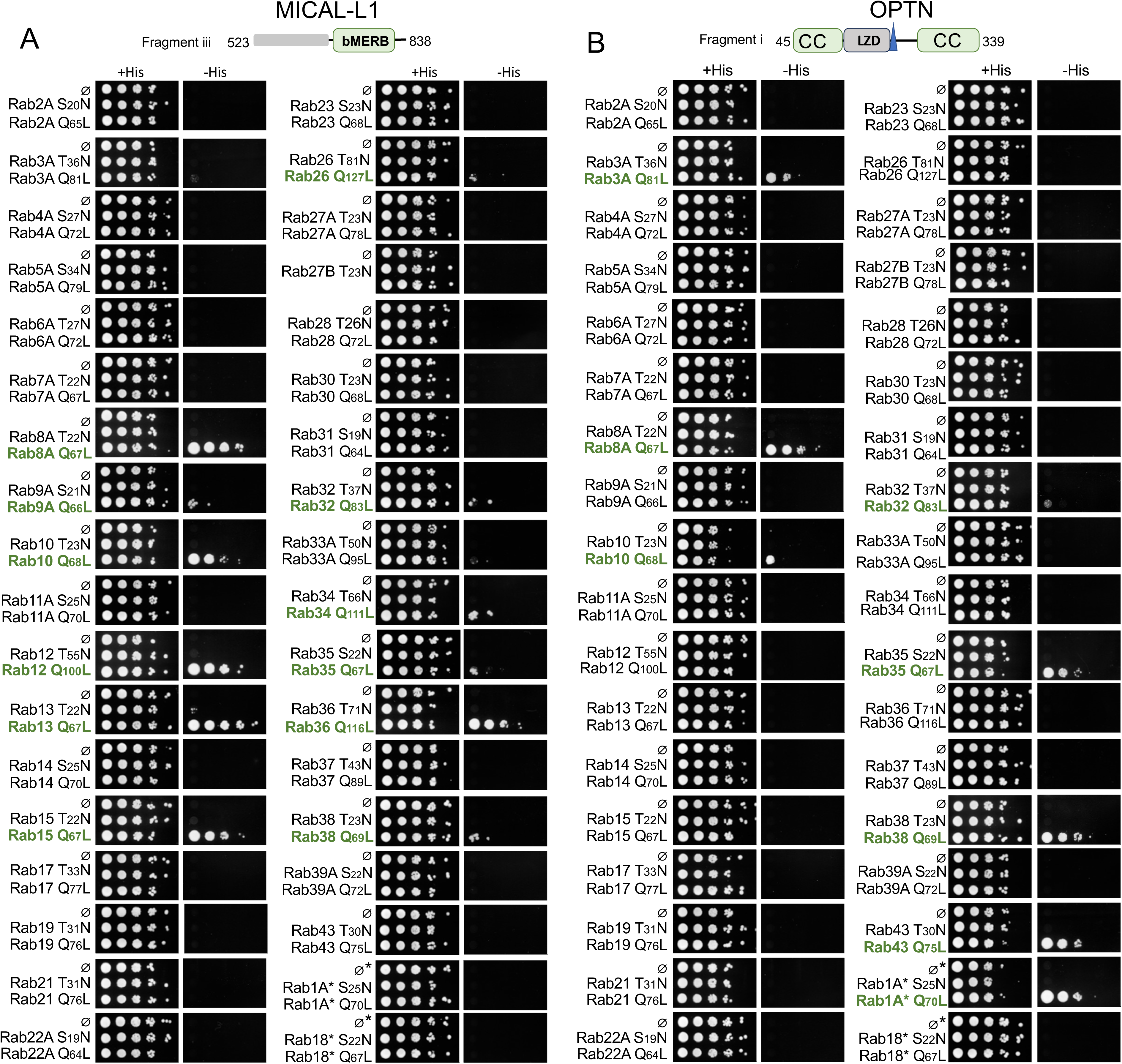
Full binary Y2H panel for MICAL-L1 and OPTN. Binary Y2H spot assays for MICAL-L1 C-terminal fragment (aa 523-838, bMERB-containing) and OPTN N-terminal fragment (aa 45-339) against the complete Rab GTPase panel in GTP-locked (Q, Q>L) and GDP-locked (T, S/T>N) conformations on permissive (+His) and selective (−His) plates. Positive interactions are highlighted green. Vector-only control is denoted (Ø).

**Supplementary Figure 3.**
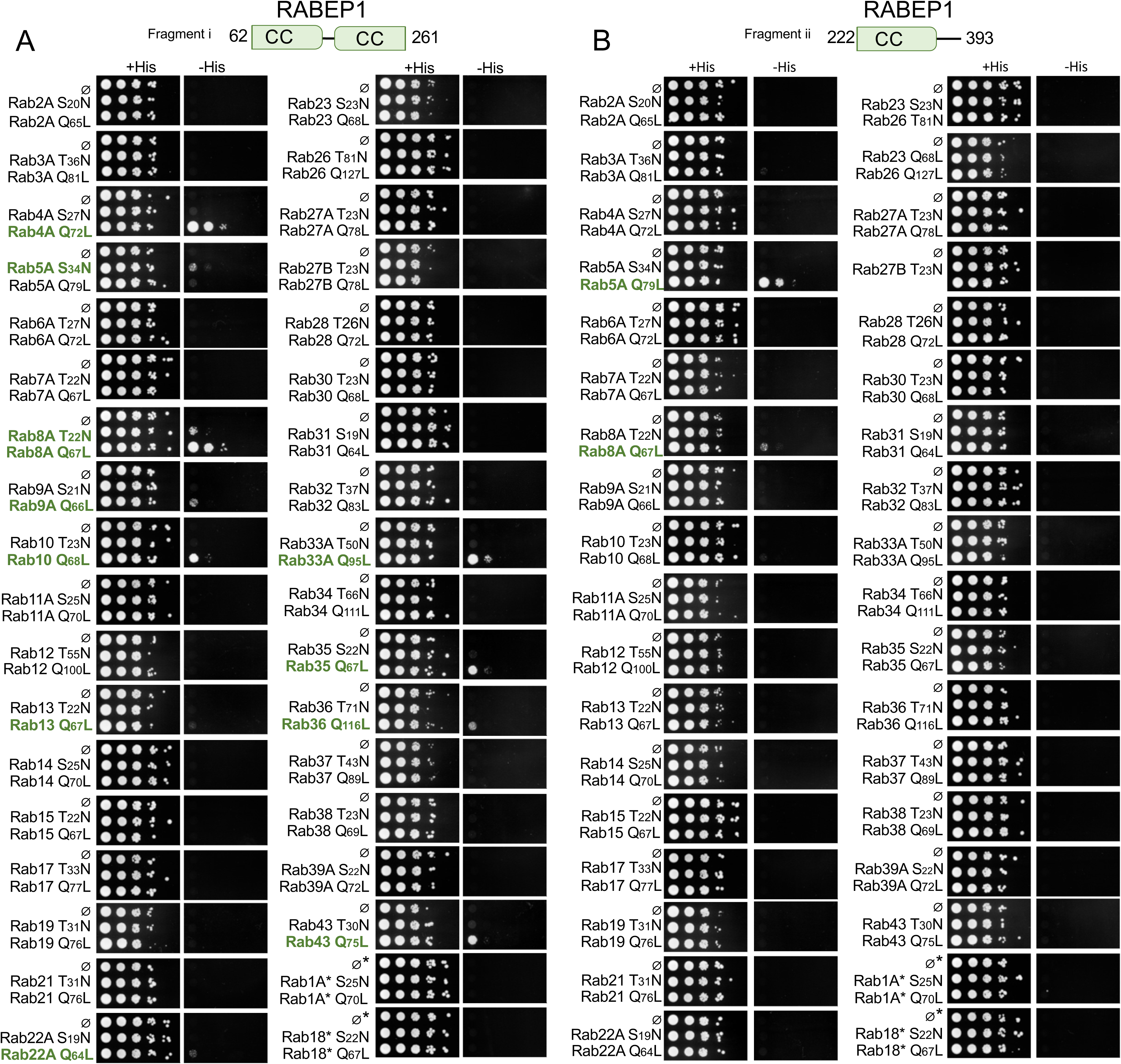
Full binary Y2H panel for RABEP1. Binary Y2H spot assays for RABEP1 fragment i (aa 62-261) and fragment ii (aa 222-393) against the complete Rab GTPase panel on permissive (+His) and selective (−His) plates, for both GTP-locked (Q>L) and GDP-locked (S/T>N) Rab baits. Fragment i shows GTP-dependent interaction with multiple Rabs; fragment ii shows selective interaction with Rab5 and weak interaction with Rab8. Positive Y2H interactions are highlighted in green. Vector-only control is denoted (Ø).

**Supplementary Figure 4.**
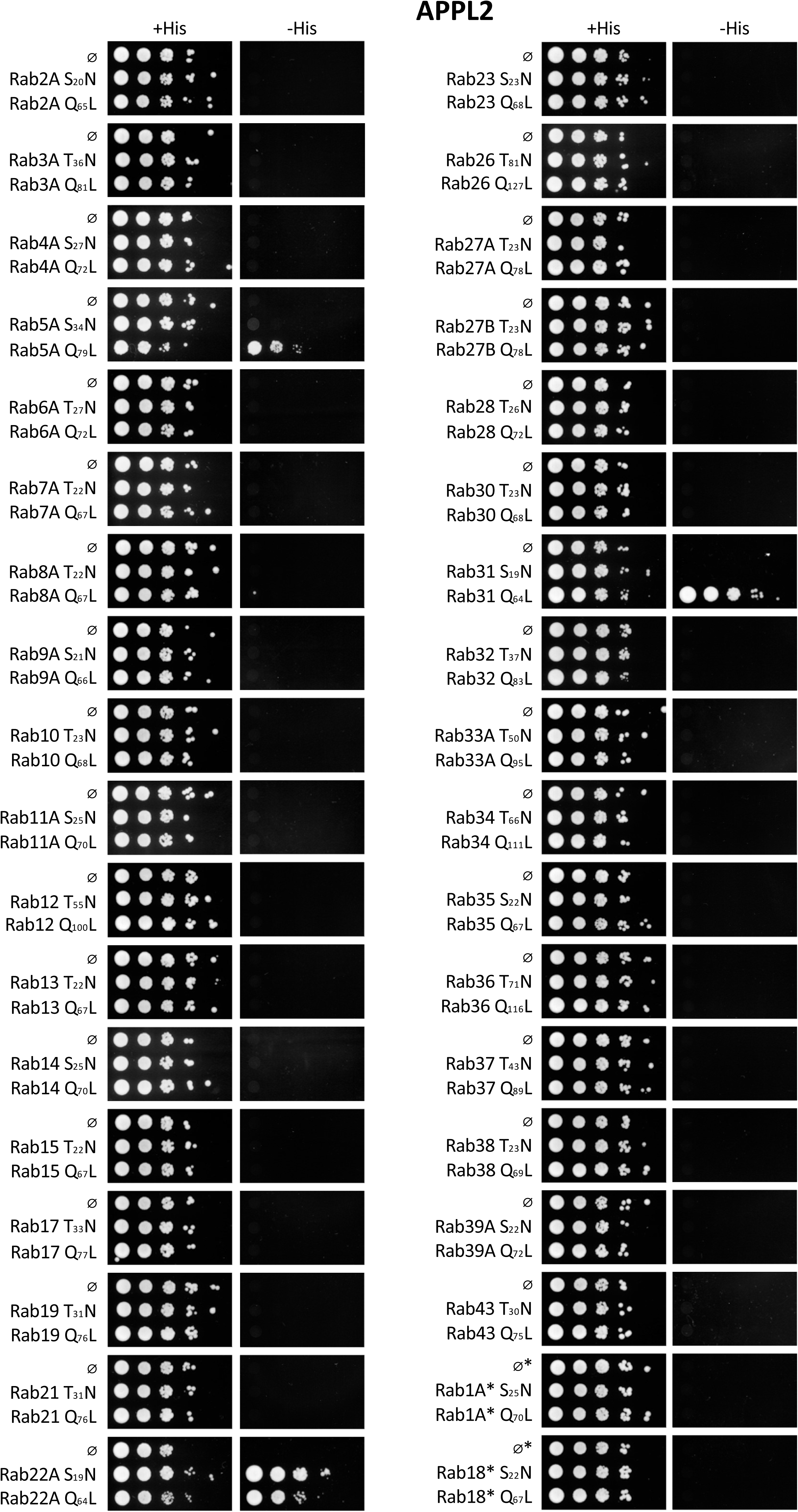
Full binary Y2H Rab panel for APPL2.

**Supplementary Figure 5.**
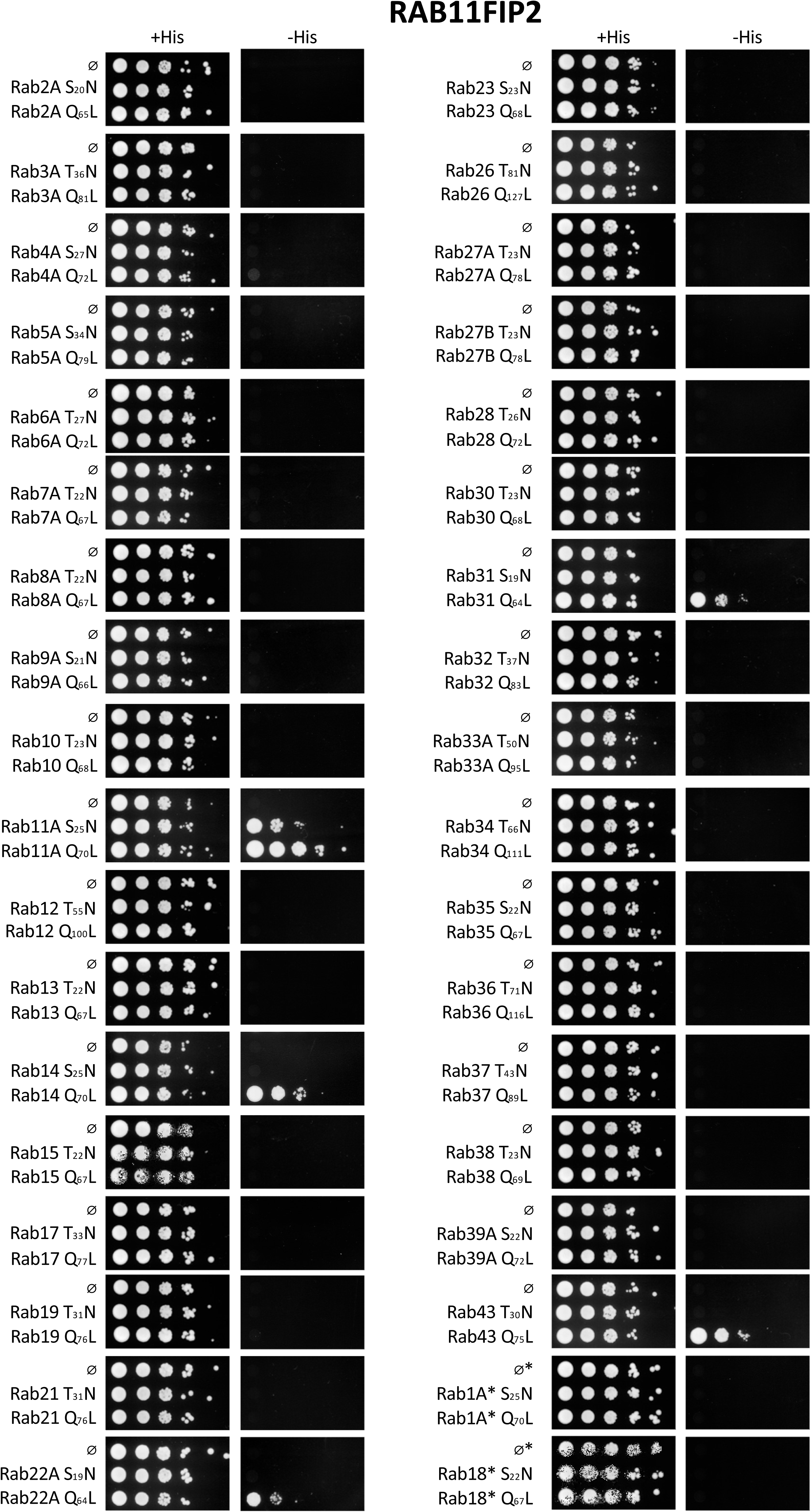
Full binary Y2H Rab panel for RAB11FIP2.

**Supplementary Figure 6.**
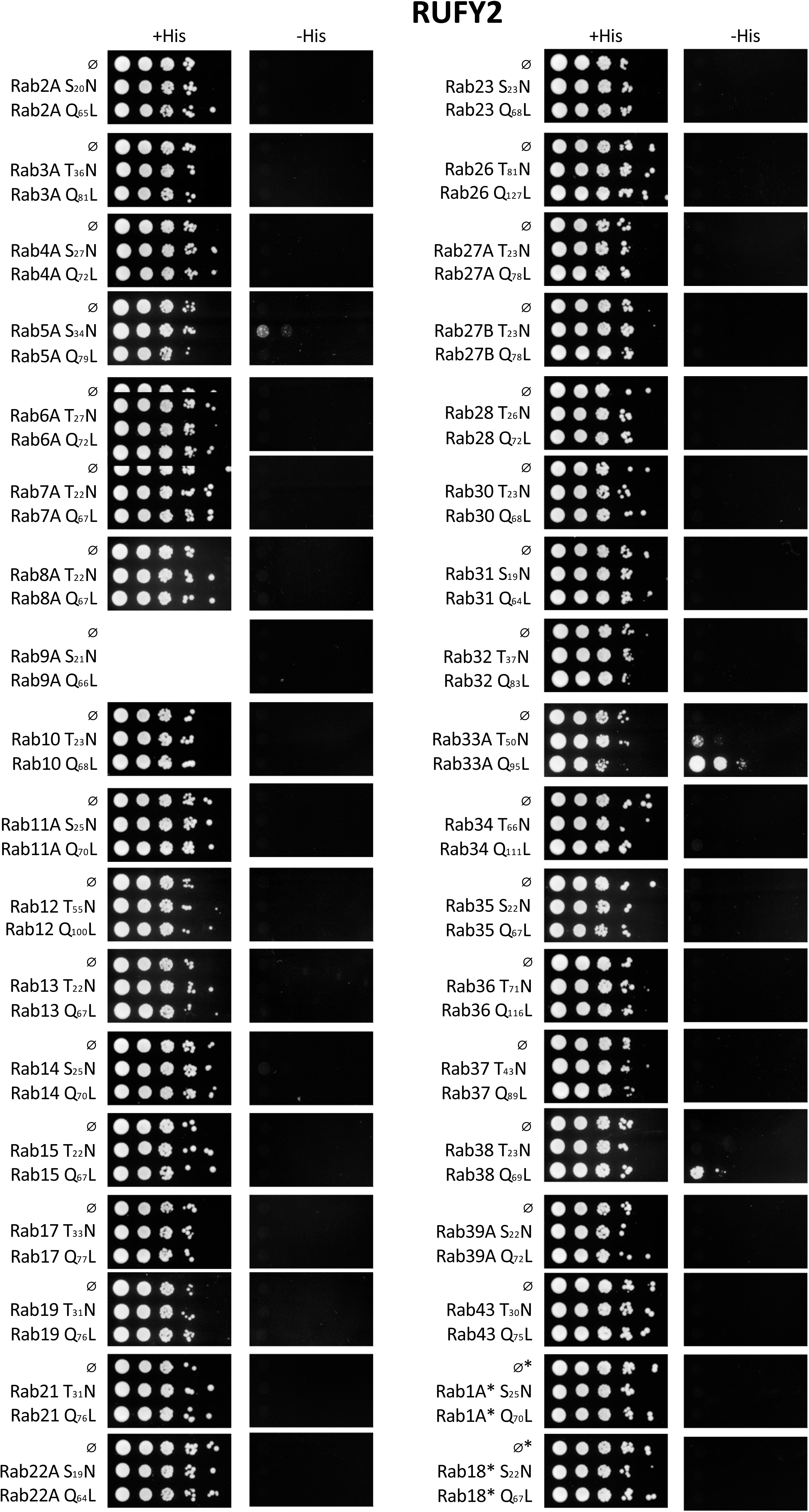
Full binary Y2H Rab panel for RUFY2.

**Supplementary Figure 7.**
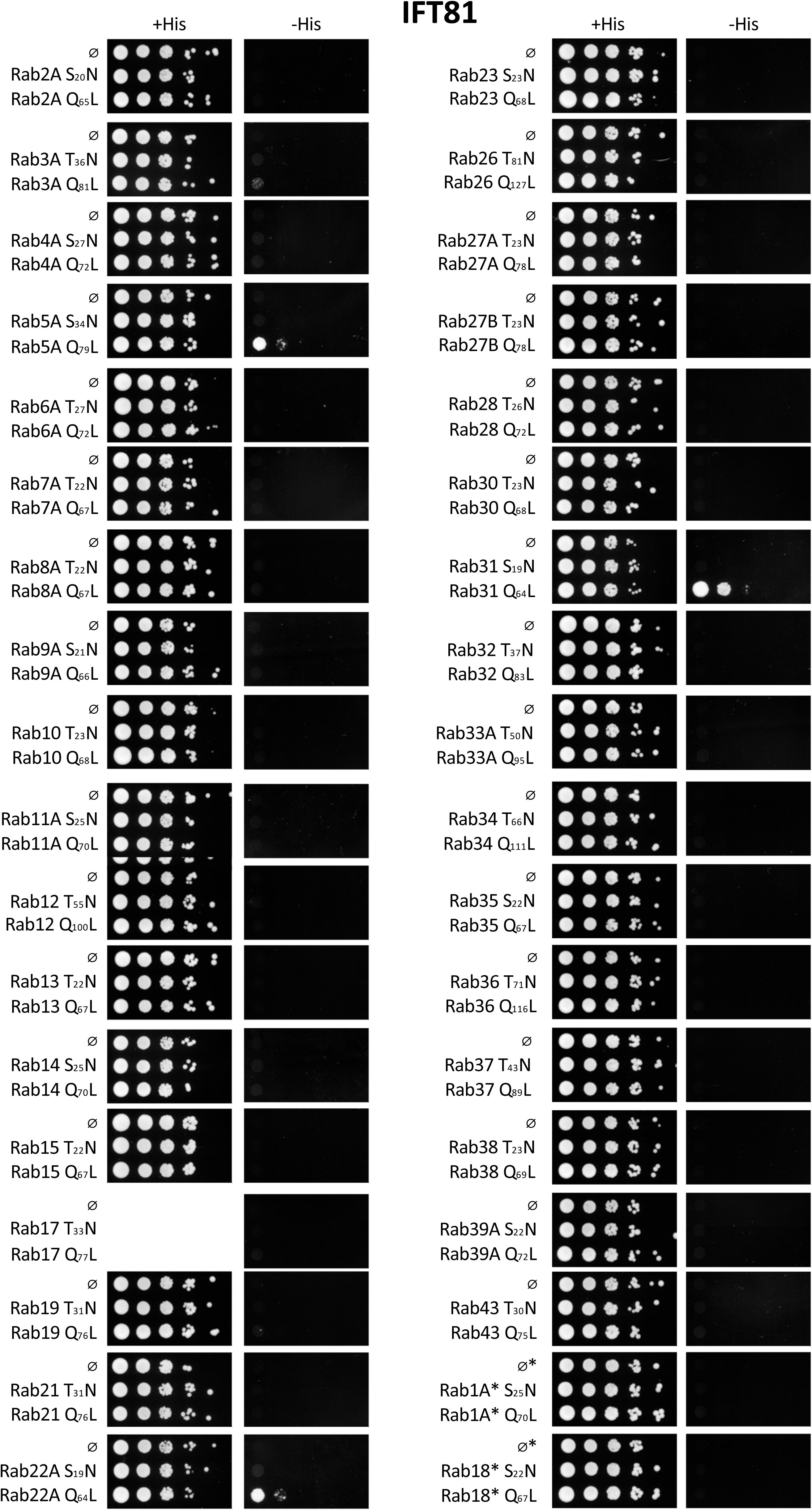
Full binary Y2H Rab panel for IFT81.

**Supplementary Figure 8.**
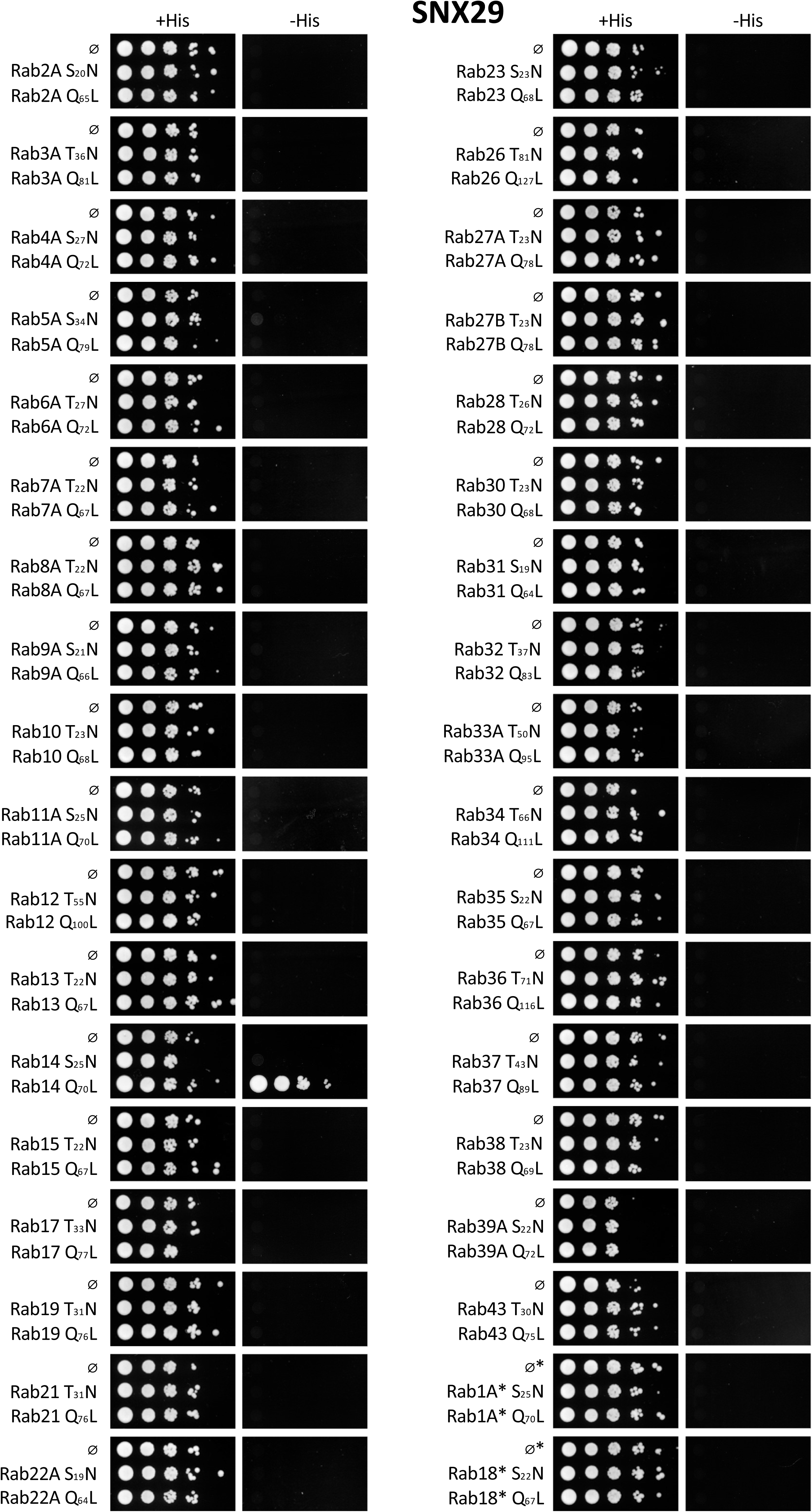
Full binary Y2H Rab panel for SNX29.

**Supplementary Figure 9.**
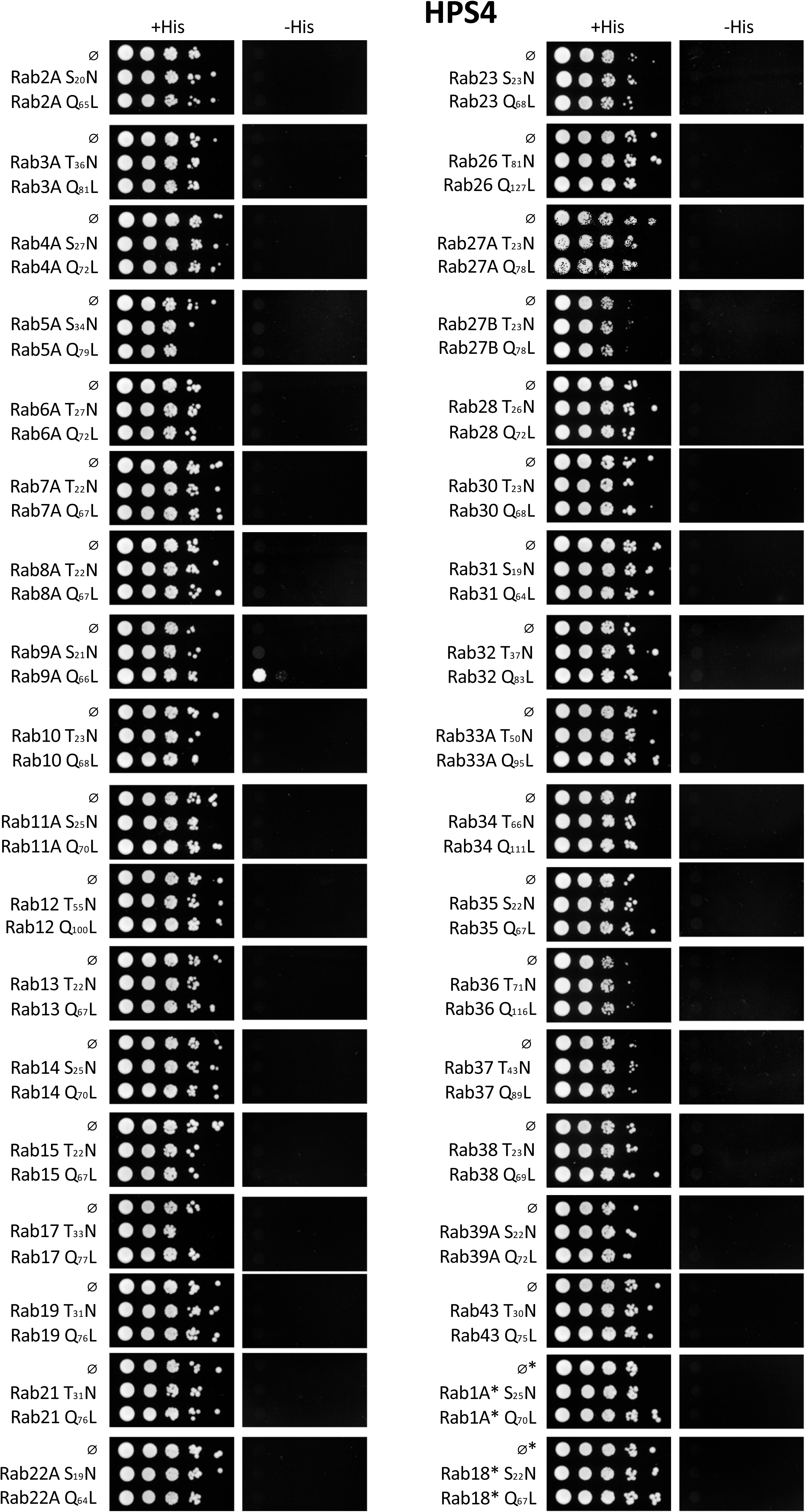
Full binary Y2H Rab panel for HPS4.

**Supplementary Figure 10.**
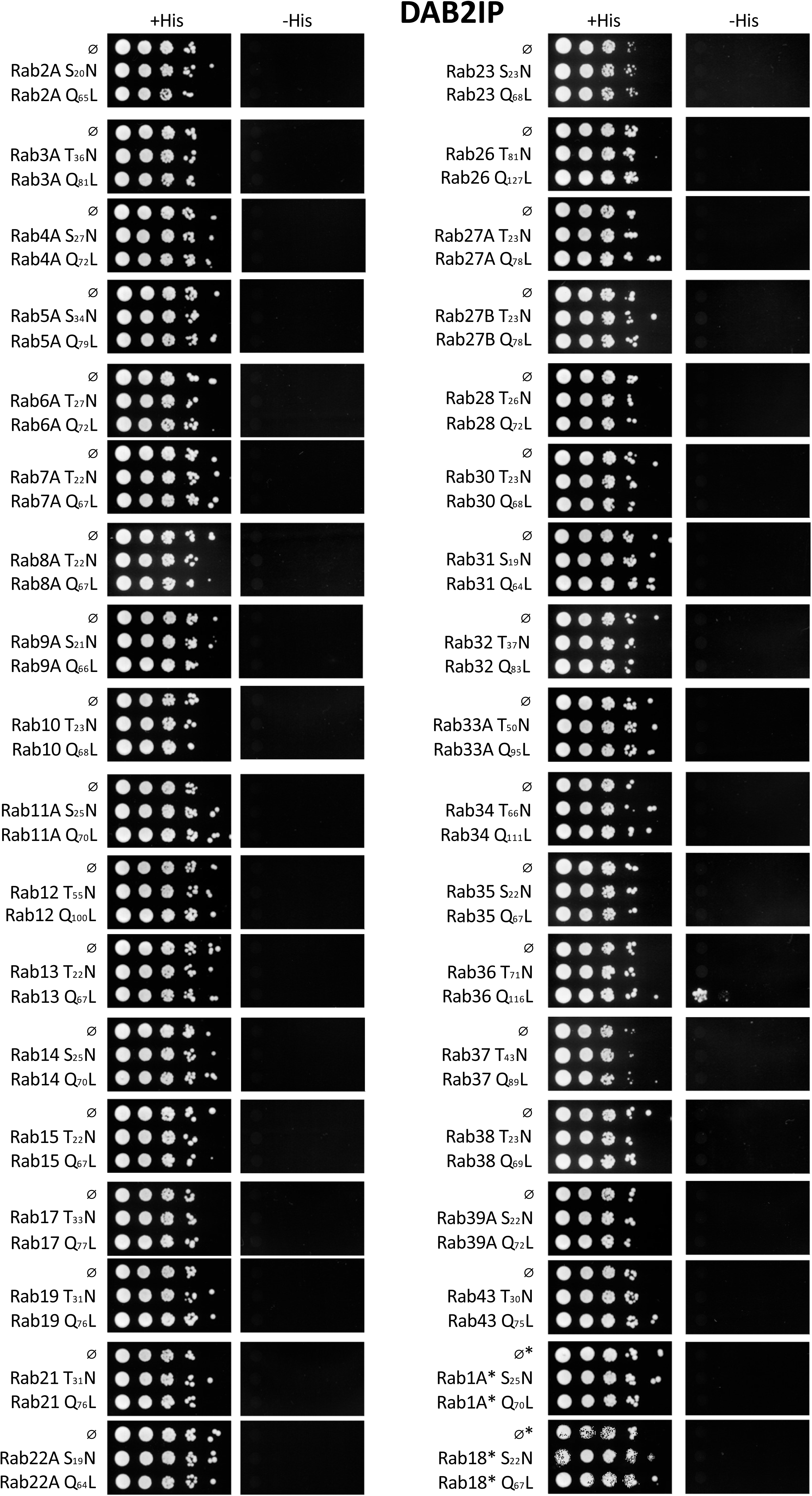
Full binary Y2H Rab panel for DAB2IP.

**Supplementary Figure 11.**
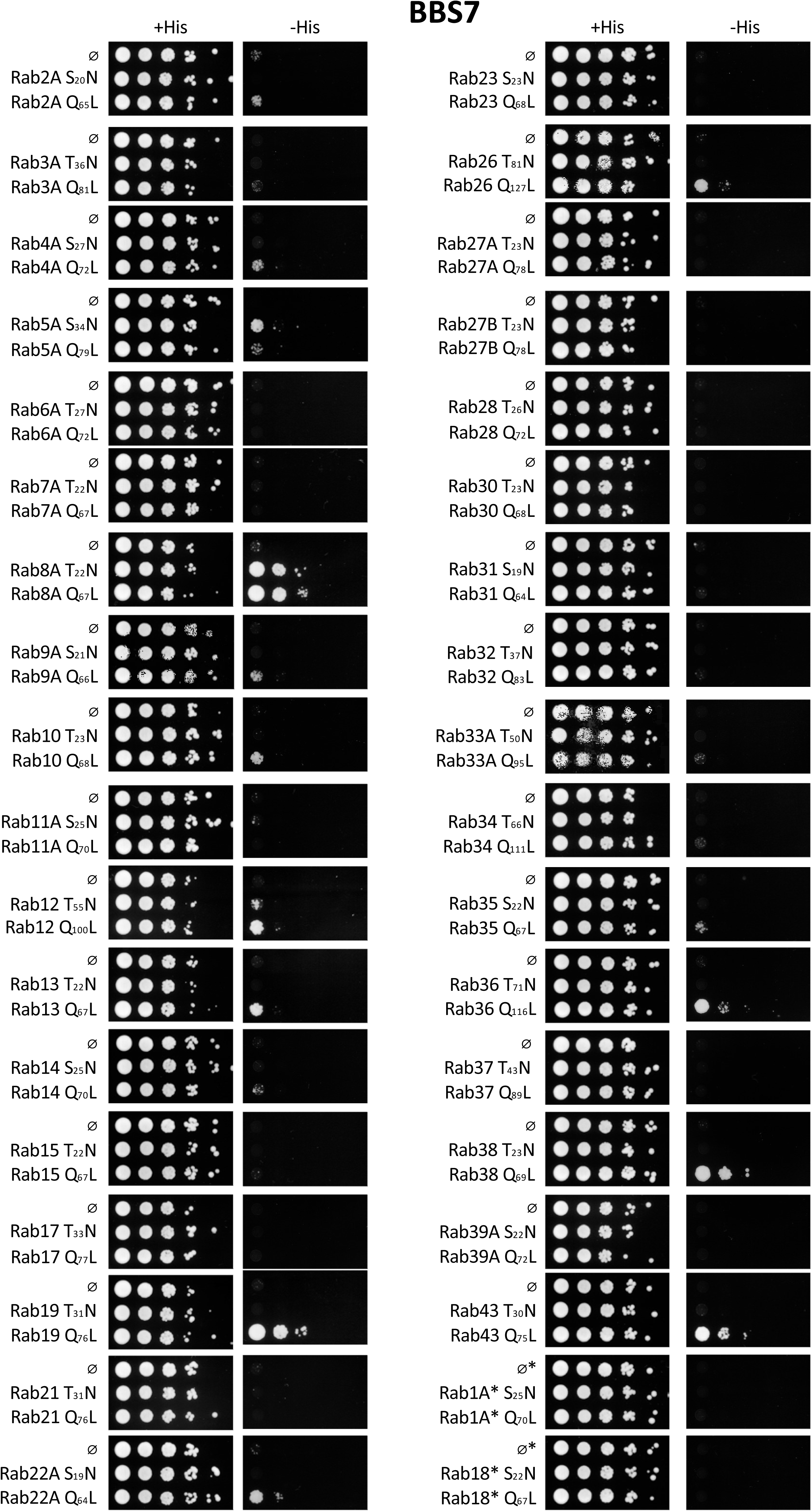
Full binary Y2H Rab panel for BBS7.

**Supplementary Figure 12.**
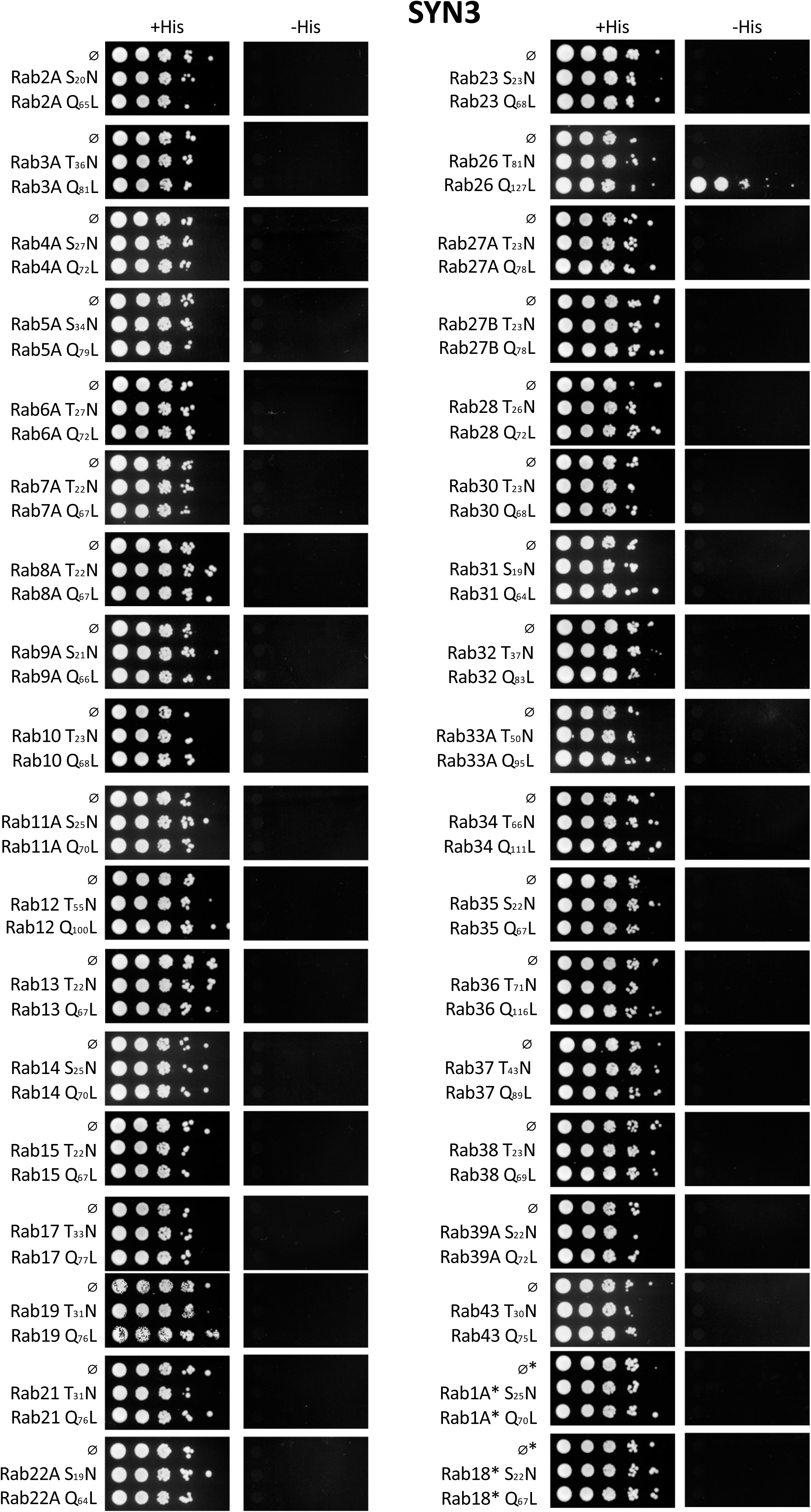
Full binary Y2H Rab panel for SYN3.

**Supplementary Figure 13.**
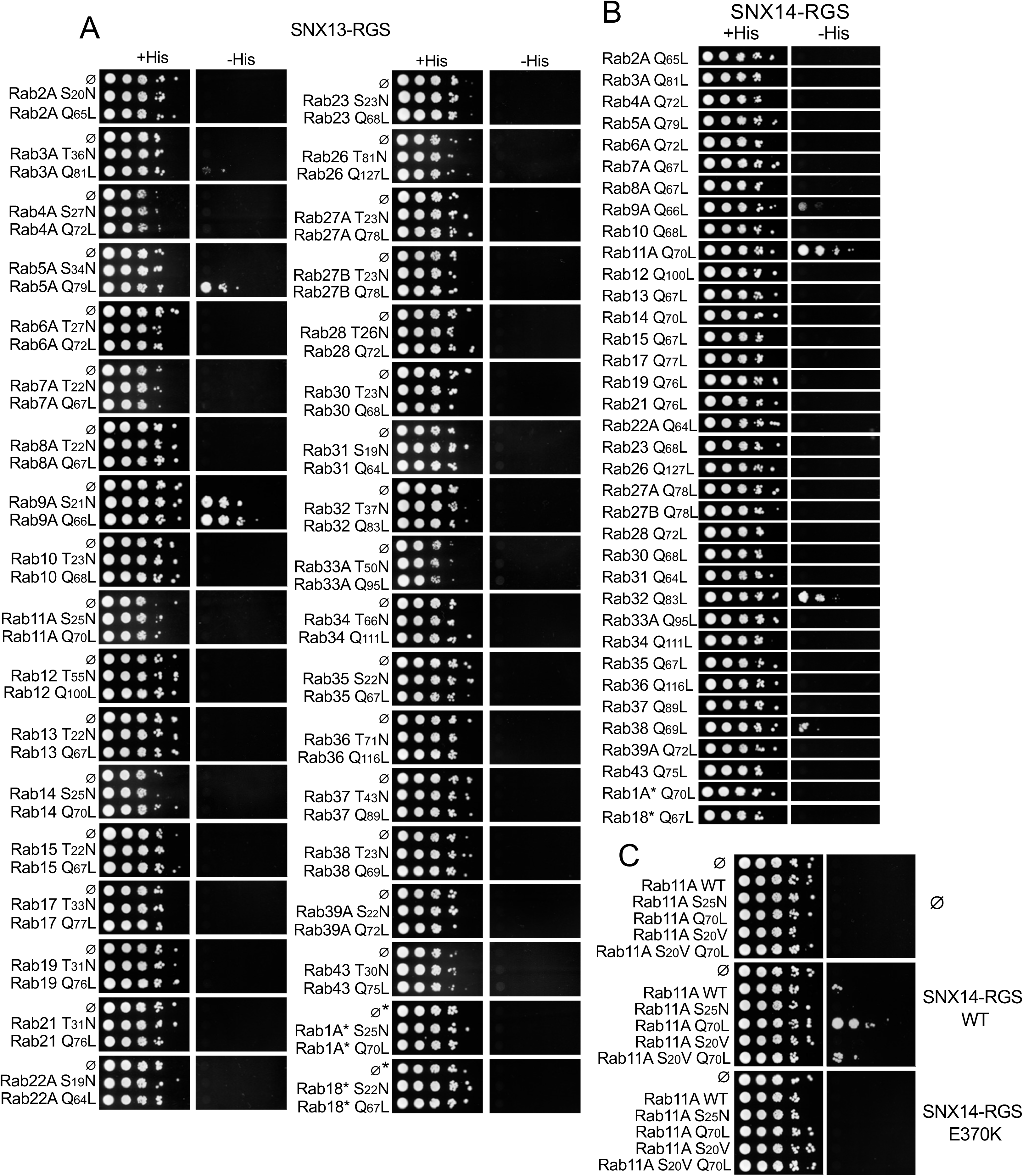
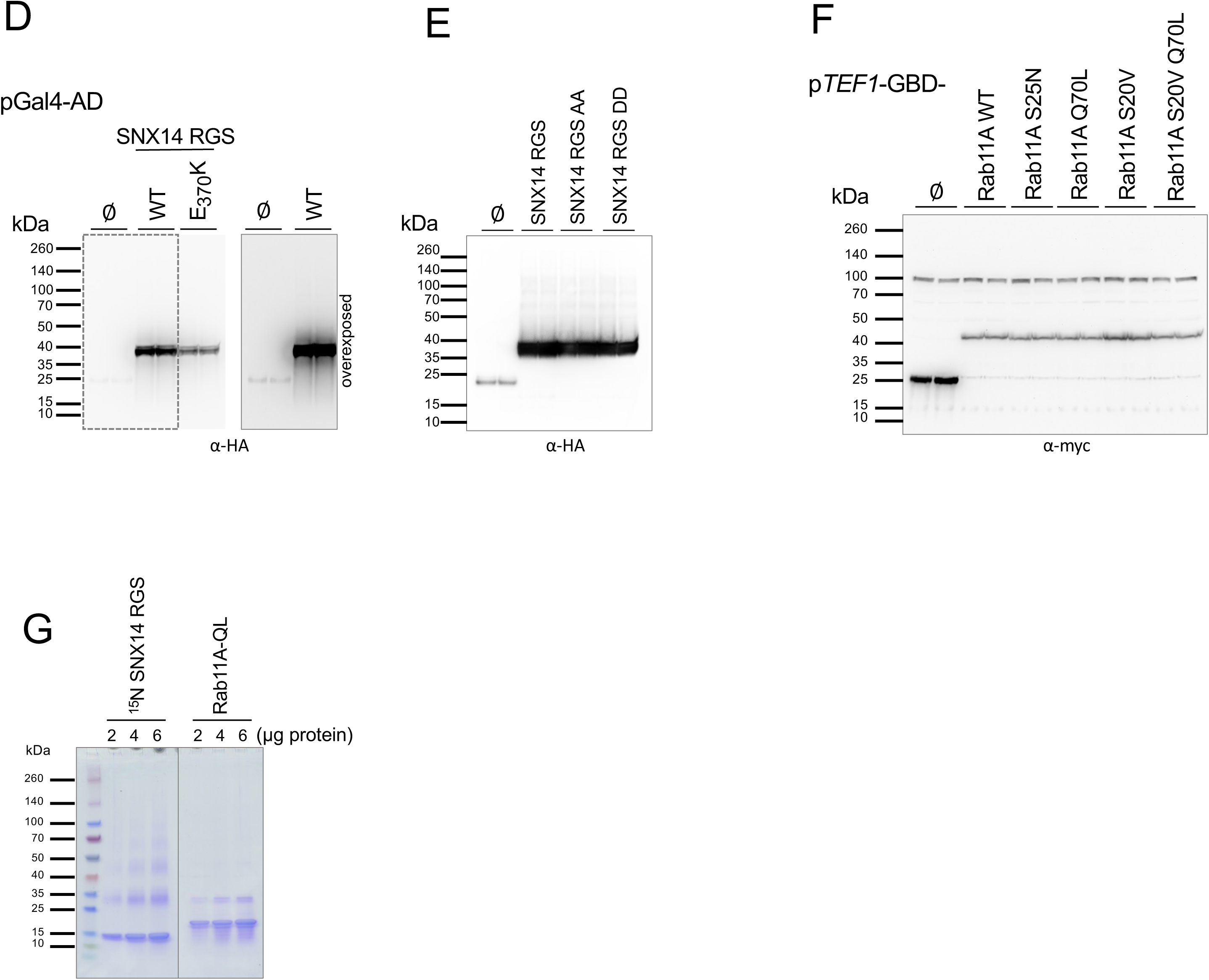
**A-C.** Vector-only control is denoted (Ø). **A.** Binary screens of the Rab panel with the SNX13-RGS domain recovered from the DEEPN screen. **B.** Binary screens of the Rab panel of the GTP-bound conformation against the SNX14 RGS domain. **C.** Y2H interaction of the indicated Rab11 mutants with vector alone, the SNX14 RGS domain, or the RGS domain with the E370K mutation. Both the S20V and the Q70L mutations are thought to lock Rab11 in a GTP conformation. However, the S20V mutation on its own blocks binding of Rab11 to SNX14 RGS domain and it diminishes binding to the Q70L mutant when in combination. **D.** Levels of the wild type and E370K mutant SNX14 RGS Gal4 activation domain fusion protein bait in duplicate transformants. **E.** Levels of the SNX14 RGS domain with phosphorylation sites mutated to Alanine or Aspartate compared to wild type SNX14 RGS. **F.** Immunoblots of the indicated Rab11 Gal4-GBD fusion proteins used in C. **G.** Pooled and dialyzed samples used for NMR analysis were analyzed by SDS-PAGE and Coomassie blue staining.

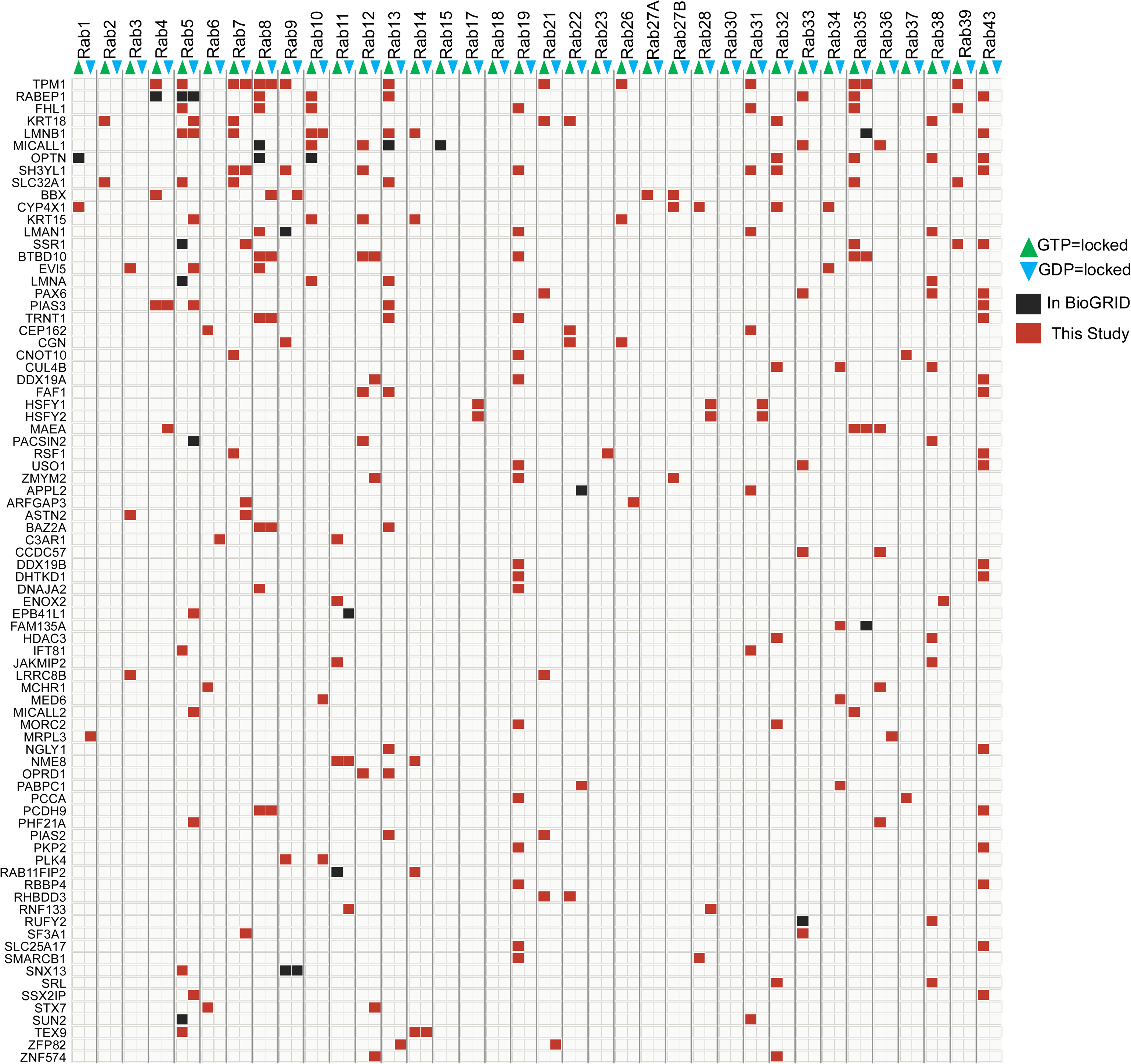

